# Dipeptidase 1 is a functional receptor for coronavirus PHEV

**DOI:** 10.1101/2025.01.09.632101

**Authors:** Jérémy Dufloo, Ignacio Fernández, Atousa Arbabian, Ahmed Haouz, Luis G. Gimenez-Lirola, Félix A. Rey, Rafael Sanjuán

## Abstract

Coronaviruses of the subgenus *Embecovirus* include several relevant pathogens such as the human seasonal coronaviruses HKU1 and OC43, bovine coronavirus, and porcine hemagglutinating encephalomyelitis virus (PHEV), among others. While sialic acid is thought to be required for embecovirus entry, protein receptors are unknown in most cases. Here we show that PHEV does not require sialic acid for entry and uses dipeptidase 1 (DPEP1) as a receptor. Cryo-electron microscopy revealed that PHEV, unlike other embecoviruses, samples open and closed conformations of its spike trimer at steady state. We found that the receptor binding domain (RBD) of the PHEV spike shares no detectable sequence homology or receptor usage with those of closely related viruses. In contrast, the X-ray structure of the RBD/DPEP1 complex showed that the elements involved in receptor binding are conserved across embecoviruses, revealing a striking versatility of the RBD to accommodate highly variable sequences that confer novel receptor specificities.

## Introduction

*Embecovirus* is one of the five subgenera of the genus *Betacoronavirus* and is characterized by encoding a hemagglutinin esterase (HE). The main embecovirus taxon is the betacoronavirus 1 species, which includes ten host-specific viruses infecting artiodactyls such as the porcine hemagglutinating encephalomyelitis virus (PHEV) and bovine coronavirus (BCoV), but also other mammals including humans, where they cause seasonal colds (OC43) or diarrhea (human enteric coronavirus strain 4408). Other relevant embecovirus species include the human common cold virus HKU1 and murine hepatitis virus (MHV). It is believed that present-day embecoviruses derive from ancestral rodent viruses that have diversified through cross-species transmission^1^. For example, it has been estimated that BCoV and OC43 diverged approximately 130 years ago following a zoonotic or reverse zoonotic event^2^.

The entry of coronaviruses into host cells is mediated by the spike protein trimer, which is responsible for binding to a specific receptor and triggering the fusion of viral and cellular membranes. The C-terminal domain of the spike S1 subunit typically acts as a protein-receptor binding domain (RBD)^3^, but the receptor it recognizes remains unknown for most embecoviruses, including all members of the betacoronavirus 1 species. For many coronaviruses, it was shown that the RBD can adopt “up” and “down” conformations that determine spike opening, which is a pre-requisite for membrane fusion. However, all embecovirus spikes, including those of MHV^4^, OC43^5^ and HKU1^6^, have been observed exclusively in closed conformation at steady state, suggesting that binding to a ligand is necessary to promote or stabilize spike opening and allow viral entry, as shown for HKU1^7^.

The N-terminal domain (NTD) of the embecovirus spike S1 subunit harbors a conserved glycan-binding site that interacts with 9-*O*-acetyl-sialic acid in HKU1, OC43, BCoV, and PHEV^8,9^, and this interaction is necessary for HKU1, OC43 and BCoV entry^5,8^. The importance of glycan attachment is further highlighted by the presence of a HE on the embecovirus envelope, which acts as a receptor-degrading enzyme analogous to the influenza C/D hemagglutinin-esterase-fusion protein^10,11^. Although this suggests that sialoglycans might function as embecovirus receptors, recent findings on HKU1 reveal a more complex scenario. HKU1 enters cells through a two-step process in which binding of the S1 NTD to 9-*O*-acetyl-sialic acids^8,12,13^ triggers spike opening^7^, allowing the RBD to interact with transmembrane protease serine 2 (TMPRSS2), which acts as a functional HKU1 receptor^14–17^. Nevertheless, whether the entry of other embecoviruses dually depends on binding to sialic acids and a cell surface protein remains unknown.

Here, we identified dipeptidase 1 (DPEP1) as a functional receptor for coronavirus PHEV. We showed that, contrary to HKU1 and other betacoronavirus 1 members, the PHEV spike does not require binding to sialic acid for entry. We thoroughly characterized DPEP1 as a sufficient PHEV receptor and obtained the X-ray structure of the PHEV RBD in complex with DPEP1. Cryo-electron microscopy (cryo-EM) of the PHEV spike trimer revealed that in contrast to other embecoviruses, it can be found in open conformations in the absence of ligand binding and that it can only bind DPEP1 in its open state. We found that RBD sequences vary extensively within the betacoronavirus 1 species and that DPEP1 usage is specific to PHEV. In contrast, the structural elements involved in receptor binding are conserved across embecoviruses. Our results highlight the diversity of embecovirus entry mechanisms and reveal that the embecovirus RBD is capable of accommodating highly divergent sequences to achieve receptor shifts.

## Results

### Sialic acid is dispensable for PHEV spike-mediated viral entry

Given the importance of sialic acid binding for embecovirus entry and the conservation of the sialoglycan binding site in the PHEV spike NTD^8^, we tested whether sialic acid was required for PHEV entry. To this aim, HCT-15 cells were pretreated with neuraminidase (NA) to remove all (α2,3)-, (α2,6)- and (α2,8)-linked sialic acids prior to infection with a vesicular stomatitis virus (VSV) pseudotype bearing the PHEV spike^18^. NA treatment only modestly decreased PHEV entry (20 ± 0.05% reduction compared to untreated cells; t-test: P = 0.02; **Figure 1A**). Similarly, a W90A PHEV spike mutant known not to interact with sialic acid^8^ showed only a twofold infectivity loss (7.22 ± 0.4% vs 3.5 ± 0.14%; t-test: P = 0.0001; **Figure 1B**). These results indicate that sialic acid may act as a PHEV attachment factor but is not strictly required for viral entry, in contrast to other betacoronavirus 1 members and HKU1.

**Figure 1.**
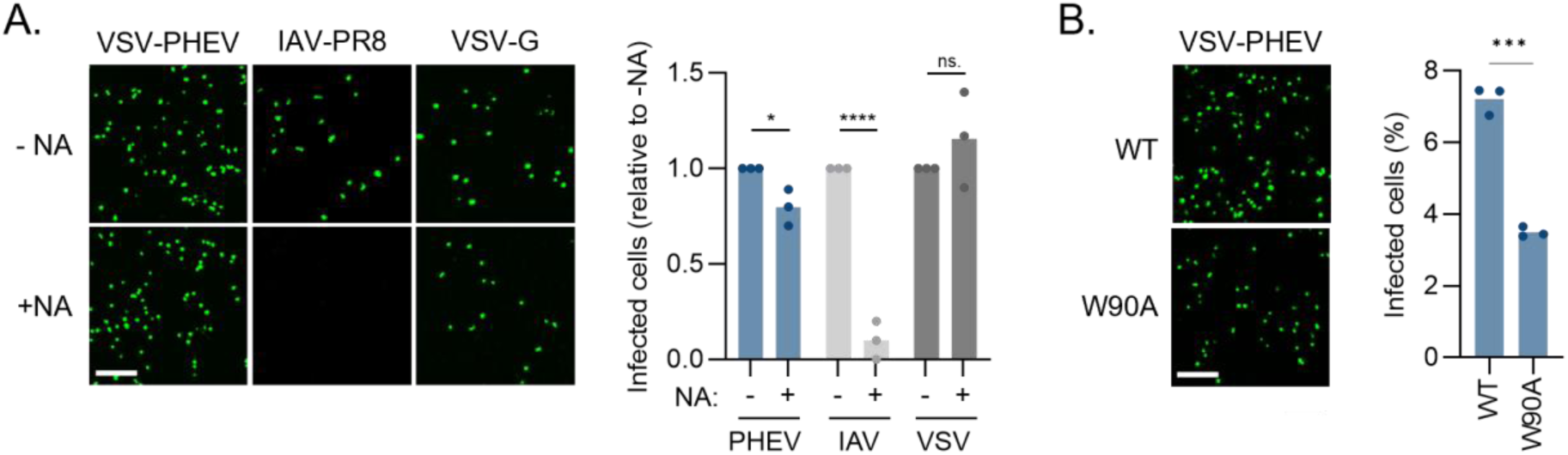
PHEV does not require sialic acid for entry. **A.** HCT-15 cells were treated or not with neuraminidase and infected with VSV-G and PHEV VSV pseudotypes or IAV (PR8-GFP). Left: representative images are shown (scale bar: 200 µm). Right: the percentage of infected cells relative to the no NA treatment is shown. Each dot represents an independent experiment (n = 3). **B.** HCT-15 cells were infected with a WT or W90A PHEV spike mutant pseudotype. Left: representative images are shown (scale bar: 200 µm). Right: the percentage of infected cells is shown. Each dot indicates a technical replicate (n = 3). The bars represent the mean and the levels of statistical significance of two-sided unpaired t-tests are shown: **** P < 0.0001, *** P < 0.001, * P < 0.05, ns: not significant.

### Identification of a PHEV candidate receptor

The above results strongly suggest that PHEV uses a protein receptor for entry. To identify potential candidates, we systematically quantified PHEV pseudotype infectivity in 48 human cell lines from the NCI-60 panel^18^. We observed infection in 16 out of 48 cell lines tested, ranging from 0.13% in T47D cells to 39% in SW-620 cells (**Figure S1A-B**). Since the NCI-60 panel has been previously characterized by transcriptomics^19,20^, we correlated PHEV pseudotype infectivity with the expression levels of 7,694 genes encoding cell surface proteins (**Figure S1C**). The expression of six genes correlated positively and significantly with infectivity after adjusting for false discovery rates (**Figure 2A**). The top hit was DPEP1 (Pearson r = 0.766, P < 0.0001; **Figure 2B**), a GPI-anchored peptidase involved in renal metabolism.

**Figure 2.**
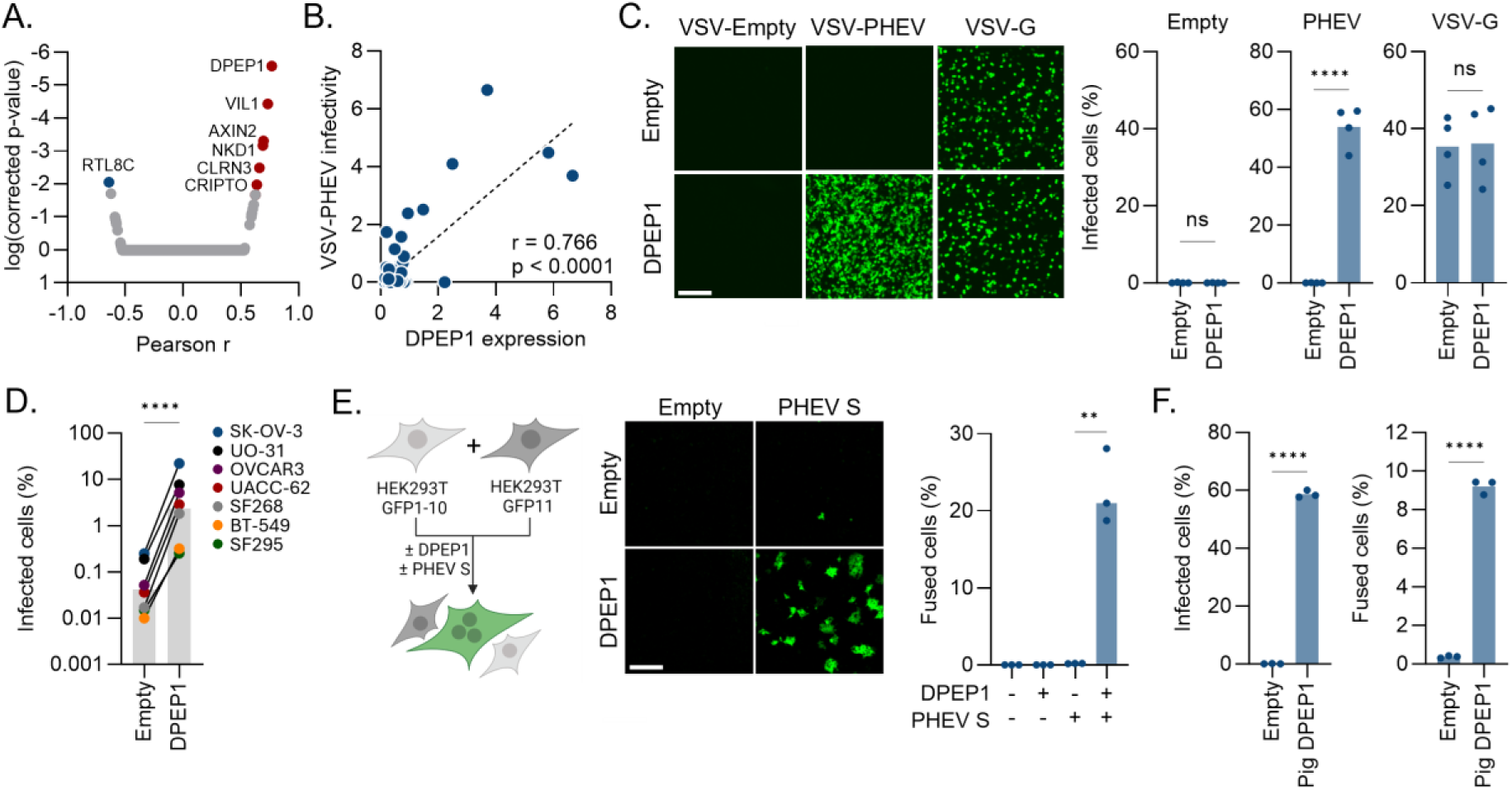
Dipeptidase 1 (DPEP1) triggers PHEV spike-mediated fusion. **A.** PHEV pseudotype infectivity in the NCI-60 panel was correlated with the expression of 7694 genes encoding membrane-associated proteins. The Pearson correlation coefficients and the Bonferroni-corrected p-values for each gene are shown. Each dot represents a different gene (n = 7694). Genes in red and blue are the ones with significant (P < 0.01) positive and negative correlation with infectivity, respectively. The names of these genes are indicated. **B.** Correlation between the expression of DPEP1 and PHEV infectivity in the NCI-60 cell lines. Each point corresponds to a different cell line (n = 47). Pearson r and p-value are indicated. **C.** HEK293T cells were transfected with a DPEP1-expressing plasmid or an empty vector and infected with empty, VSV-G and PHEV VSV pseudotypes. Cells were imaged 20 hours post-infection. Representative images are shown on the left (scale bar: 200 µm). On the right, the percentage of infected cells is shown. Each dot represents an independent experiment (n = 4). **D.** Different cell lines were transfected with a DPEP1-expressing plasmid or an empty vector and infected with PHEV pseudotypes. The percentage of infected cells at 20 hours post-infection is shown. Each dot represents a different cell line (n = 7). The bar represents the mean and the level of significance of a two-sided paired t-test is shown: **** P < 0.0001. **E.** Left panel: schematic of the cell-cell fusion assay. Briefly, HEK293T GFP-Split cells were mixed and transfected with DPEP1 and the PHEV spike (or empty vectors). GFP-positive syncytia were quantified 24 h post-transfection. Middle panel: representative images of the cell-cell fusion assay upon DPEP1 and/or PHEV spike transfection (scale bar: 200 µm). Right panel: percentage of cell-cell fusion in cells overexpressing the PHEV spike and DPEP1 or an empty control. Each dot represents a technical replicate (n = 3). **F.** Left panel: HEK293T cells were transfected with a pig DPEP1-expressing plasmid or an empty vector and infected with a PHEV pseudotype. The percentage of infected cells is shown. Each dot represents an independent experiment (n = 3). Right panel: HEK293T GFP-Split cells were transfected with the PHEV spike and pig DPEP1 or an empty vector. The percentage of cell-cell fusion is shown. Each dot represents a technical replicate (n = 3). In **C**, **E** and **F**, the bar represents the mean and the levels of statistical significance of two-sided unpaired t-tests are shown: **** P < 0.0001, ** P < 0.01, ns: not significant.

### DPEP1 induces PHEV spike-mediated fusion

We next examined the effect of DPEP1 transfection on PHEV pseudotype infectivity in HEK293T cells. DPEP1 overexpression increased infection rates from undetectable levels (0.07 ± 0.04%) to 54 ± 3.6% (t-test: P < 0.0001; **Figure 2C**). This effect was specific to PHEV, as infection with an empty or VSV-G pseudotype was unaffected by DPEP1 overexpression (**Figure 2C**). To ensure that these results were independent of the cellular context, we tested seven additional cell lines, where we observed an average 73-fold increase in PHEV pseudotype infection upon DPEP1 overexpression (**Figure 2D**). We also used a GFP complementation assay to assess whether expression of the spike at the cell surface triggered syncytia formation (**Figure 2E**). Transfection of DPEP1 or the PHEV spike alone did not induce significant cell-cell fusion, but co-expression of both allowed strong syncytia formation (0.18 ± 0.01% in PHEV spike-transfected cells vs 22.6 ± 2.8% in PHEV spike-DPEP1 co-transfected cells; t-test: P = 0.0013). Since PHEV is a porcine virus, we confirmed that the *Sus scrofa* DPEP1 ortholog also allowed PHEV pseudotype infection and spike-mediated cell-cell fusion (**Figure 2F**). Overall, these experiments demonstrate that the porcine and human versions of DPEP1 sensitize non-susceptible cells to PHEV spike-mediated entry and induce spike-mediated cell-cell fusion.

### DPEP1 binds the PHEV spike

Since DPEP1 is a peptidase, we first sought to determine whether it acted as a true PHEV receptor or whether it promoted entry by priming the spike. To this end, we generated the catalytically inactive mutant E141D^21^. This mutant was still capable of mediating viral entry into transfected HEK293T cells and induced PHEV spike-mediated cell-cell fusion, showing that the enzymatic activity of DPEP1 is dispensable to promote PHEV entry (**Figure 3A**). We then examined whether DPEP1 bound the PHEV spike. For this, we incubated PHEV spike-transfected HEK293T cells with increasing concentrations of soluble recombinant human DPEP1 and measured DPEP1 binding by flow cytometry (**Figure 3B**). We observed a strong dose-dependent binding of soluble human DPEP1 to spike-expressing cells, which was not observed in control cells. We then expressed the PHEV spike RBD (residues 327-605) to measure its affinity for the porcine DPEP1 ectodomain by bio-layer interferometry (BLI; **Figure 3C**). The immobilized PHEV RBD interacted with DPEP1 but not with ACE2. In contrast, the SARS-CoV-2 RBD (residues 331-528, Wuhan strain) bound ACE2 but not DPEP1, confirming the specificity of the interaction (**Figure 3C**). To measure the binding affinity avoiding avidity effects, we changed the experimental set-up, immobilizing DPEP1 and using a range of PHEV RBD concentrations, determining an equilibrium dissociation constant (*K_d_*) of 1.50 ± 0.31 μM (**Figure 3D-E**). We also expressed a recombinant PHEV spike ectodomain (residues 15-1274) to confirm its interaction with DPEP1 by BLI (**Figure 3F**). Given that DPEP1 is a dimer and the spike protein is a trimer, we did not derive kinetic and affinity parameters, but the curves clearly demonstrate their interaction. Finally, as another approach to detect PHEV spike-DPEP1 interaction, we performed a competition assay using PHEV pseudotypes and soluble recombinant human DPEP1. PHEV pseudotypes were incubated with increasing amounts of soluble DPEP1 prior to infection of susceptible cells (HCT-15 or DPEP1-transfected HEK293T cells; **Figure 3G**). Soluble DPEP1 strongly inhibited PHEV pseudotype infection in both cell lines (IC_50_: 0.03 µg/mL in HCT-15 and 0.04 µg/mL in HEK293T-DPEP1 cells). In conclusion, we show that the human and porcine orthologs of DPEP1 specifically interact with the PHEV spike, further supporting its role as a functional PHEV receptor.

**Figure 3.**
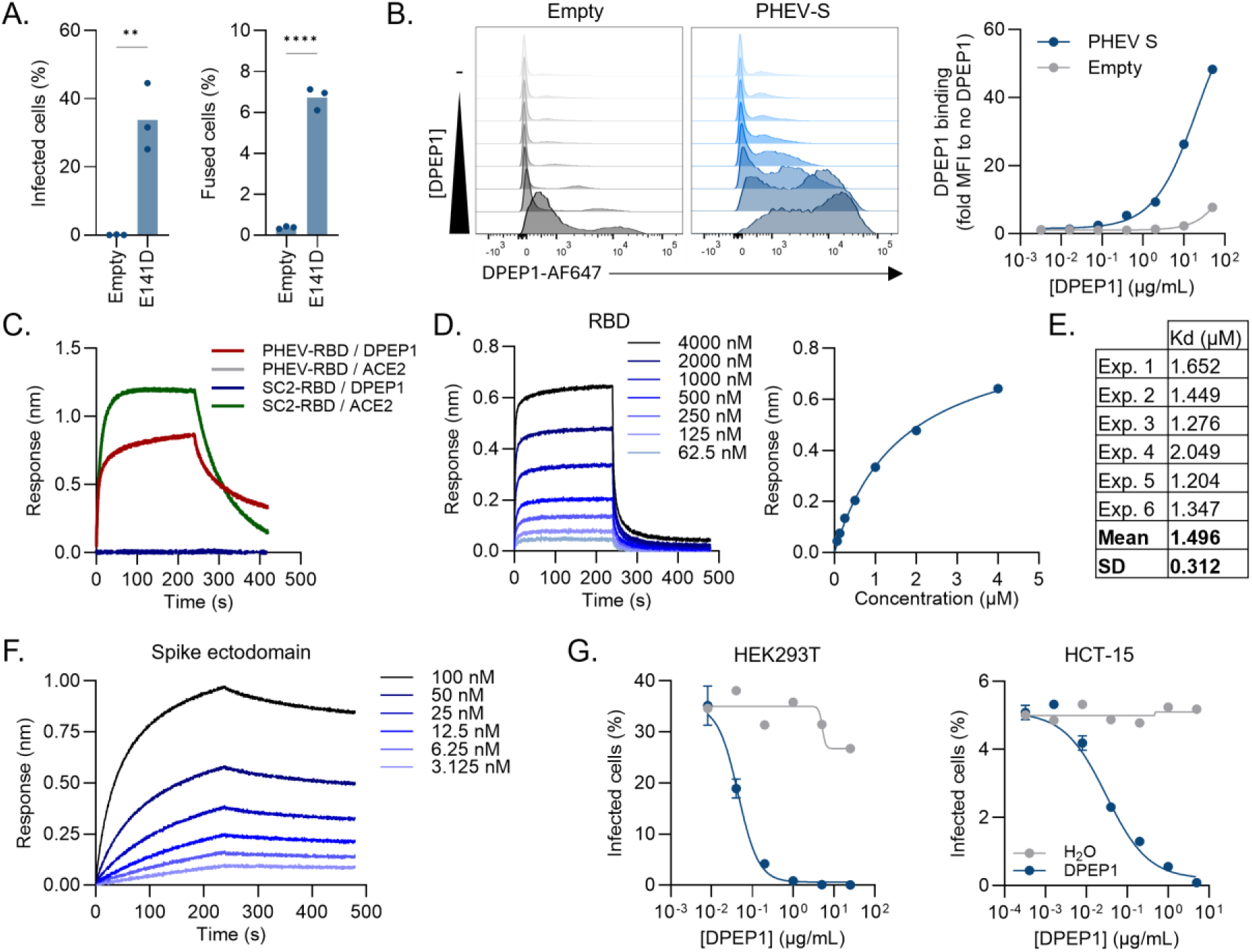
DPEP1 binds the PHEV spike. **A.** Left panel: HEK293T cells were transfected with the human DPEP1 E141D mutant or an empty vector and infected with a PHEV pseudotype. The percentage of infected cells is shown. Each dot represents an independent experiment (n = 3). Right panel: HEK293T GFP-Split cells were transfected with the PHEV spike and pig DPEP1 or an empty vector. The percentage of cell-cell fusion is shown. Each dot represents a technical replicate (n = 3). The statistical significance of a two-sided unpaired t-test is shown: ** P < 0.01, **** P < 0.0001. **B.** Flow cytometry measurement of soluble human DPEP1 binding to HEK293T cells transfected with the PHEV spike or an empty vector. Left: flow cytometry histogram of DPEP1 binding in cells transfected with the PHEV spike or an empty vector. From top to bottom, cells were incubated without (-) and with increasing amounts of soluble DPEP1. Right: the data shows the binding median fluorescence intensity (MFI) measured at each concentration of soluble DPEP1 normalized to the MFI in the absence of DPEP1, in cells transfected with an empty vector or the PHEV spike. Results from a representative experiment are shown. **C.** Binding of porcine DPEP1 or human ACE2 to PHEV or SARS-CoV-2 (SC2) RBD-coated sensors was quantified by bio-layer interferometry (BLI). One representative experiment of two is shown. **D.** Left: binding of porcine DPEP1 to PHEV RBD measured by BLI. DPE1 was immobilized on streptavidin sensors that were incubated with different RBD concentrations. Right: determination of the affinity of PHEV RBD for DPEP1 using the steady state method. The circles are experimental values and the line is the fitting of the experimental data. One representative experiment of six is shown. **E.** Affinity constants (*K_d_*) of the PHEV RBD-DPEP1 interaction obtained in the six independent experiments. The mean *K_d_* and its standard deviation (SD) are also indicated. **F.** Binding of immobilized porcine DPEP1 to the PHEV spike ectodomain measured by BLI using different spike concentrations. One representative experiment of two is shown. **G.** PHEV pseudotypes were incubated with the indicated concentration of soluble human DPEP1 (blue curve) or vehicle (grey curve, H2O) before infecting HCT-15 (left panel) or DPEP1-transfected HEK293T cells (right panel). Data show the percentage of infected cells at each soluble DPEP1 concentration (mean ± SEM of two technical replicates).

### Soluble DPEP1 interferes with full PHEV infection

Although PHEV propagation in cell culture is challenging, we set out to investigate the role of DPEP1 using the authentic virus. We thus ran the competition assay described above with PHEV strain 67N (**Figure 4A**). We incubated the virus with soluble human DPEP1 prior to inoculation of two different susceptible cell types (PK-15 cells and swine primary kidney cells, SPKC), and followed the release of viral particles into the supernatant over time by RT-qPCR. In PK-15 cells, there was significantly less virus in the supernatant at 24 and 48 h post-inoculation when the virus was preincubated with soluble DPEP1 (24 h: 2.3-fold, t-test: P = 0.0053; 48 h: 2.9-fold, P = 0.0031; **Figure 4B**). In SPKC cells, DPEP1 pretreatment significantly reduced virus release at all time points by a factor ranging from 2.5-to 3.5-fold (t-tests: P < 0.05). These results therefore confirm the role of DPEP1 in PHEV infection.

**Figure 4.**
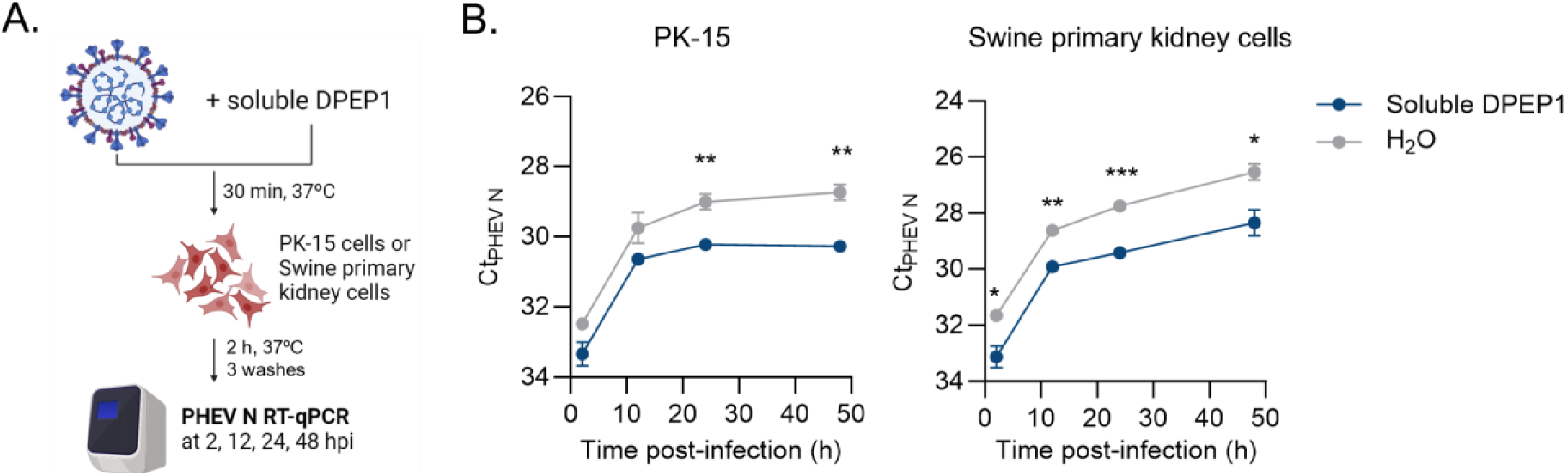
Soluble DPEP1 inhibits PHEV replication. **A.** Schematic of the full virus competition assay. Briefly, PHEV was pre-incubated with soluble human DPEP1 and used to infect PK-15 or swine primary kidney cells. Viral input was removed, cells were thoroughly washed with PBS, and viral release was followed by RT-qPCR of the supernatants at several times post-infection. **B.** PHEV N Ct values at different times post-infection with (blue) or without (grey) soluble human DPEP1. To display results in a more comprehensible way, the y-axis is reversed. Each dot represents the mean ± SEM of three replicates. The Ct values in both treatments (Soluble DPEP1 vs H_2_O) were compared at each time point using a two-sided unpaired t-test: * P < 0.05; ** P < 0.01; *** P < 0.001.

### TMPRSS2 enhances DPEP1-mediated entry but is not a receptor for PHEV

The only other protein described to bind an embecovirus RBD is TMPRSS2, which serves as a functional receptor for HKU1^14^. Moreover, TMPRSS2 activates the spike and enhances the entry of other coronaviruses (e.g. SARS-CoV-2^22^ or MERS-CoV^23^). Therefore, we investigated its role in PHEV entry. We found that the expression of TMPRSS2 also correlated with PHEV pseudotype infectivity, albeit less strongly than DPEP1 (Pearson r = 0.47, P = 0.0009; **Figures S1C and S2A**). Overexpression of TMPRSS2 alone was not sufficient to promote PHEV pseudotype infection in three non-susceptible cell lines (HEK293T, SK-OV-3 and SF-268; **Figure S2B**). However, co-expression of DPEP1 and TMPRSS2 increased viral entry in all three cell lines compared to DPEP1 alone (3.6-, 1.6-, and 1.9-fold increase in HEK293T, SK-OV-3 and SF-268 cells, respectively; **Figure S2B**). Co-expression of DPEP1 and TMPRSS2 also resulted in a 60% increase in spike-mediated cell-cell fusion compared to when DPEP1 was expressed alone (**Figure S2C**). These results indicate that TMPRSS2 enhances DPEP1-dependent entry but is unlikely to be a functional receptor for PHEV, thus indicating that receptor usage is not conserved between the HKU1 and PHEV embecoviruses.

### DPEP1 is a PHEV-specific receptor

Given that no betacoronavirus 1 receptor has been identified to date, we set out to determine whether DPEP1 was a general protein receptor for the species. To test this, we measured the binding of recombinant human DPEP1 to cells overexpressing various betacoronavirus 1 spikes (i.e. OC43, BCoV, CRCoV, ECoV). The HKU24 spike was included as a non-betacoronavirus 1 embecovirus. Despite the detectable expression of all spikes, DPEP1 binding could only be measured in PHEV-transfected cells, suggesting that DPEP1 is a PHEV-specific receptor (**Figure S3A**). To understand whether potential spike diversity might explain this receptor usage specificity, we built a phylogenetic tree of all embecovirus spike proteins (**Figure S3B**). Although betacoronavirus 1 spikes form a monophyletic group, PHEV and equine coronavirus (ECoV) differed from the rest of betacoronavirus 1 members. A region of about 100 residues in their RBD was indeed highly divergent from all other betacoronavirus 1 spikes and from each other (residues 457-559 of the PHEV spike; **Figure S3C**). This region of the PHEV RBD exhibited no significant similarity with any other viral sequence deposited in GenBank and the structure-based remote homology detection tool pLM-Blast^24^ showed no obvious hits.

### X-ray structure of the PHEV RBD in complex with DPEP1

We attempted to predict the structure of the DPEP1/PHEV RBD complex using AlphaFold3^25^, but the resulting interface confidence metrics for the predicted complex were very low (ipTM: <0.3). We therefore crystallized the PHEV RBD in complex with porcine DPEP1 and determined the X-ray structure at 2.25 Å resolution (the X-ray diffraction data collection and model refinement statistics are listed in **Table S1**). The asymmetric unit of the crystal contained one DPEP1 dimer with two RBDs bound (**Figure 5A**). Porcine DPEP1 has an α/β barrel architecture that conserves the secondary structure elements, overall fold (root mean square deviation, RMSD: 0.931 Å for 5446 atoms) and glycosylation sites (N57 and N279) of the human ortholog crystallized previously (PDB: 1ITQ)^26^. The active site has two zinc ions, one coordinated by H36, D38 and E141 and the second by H214, H235 and E141. The site of interaction with the RBD is on the face opposite to the active site and is formed by helices α_a_, α_2a_ and α_e_ (**Figure 5B**, **Figure S4**), in line with the catalytically inactive DPEP1 variant (E141D) being able to promote pseudotype entry and cell-cell fusion. The PHEV receptor binding site is in the RBD distal region, which has the shape of an open mouth (**Figure 5B**). The upper jaw (jaw 1, j1) is formed by a long loop (residues 506-518) and the lower jaw (jaw 2, j2) consists of the short helix α9 (residues 531-533, **Figure S5**) in the highly variable region observed in betacoronavirus 1 RBDs (**Figure S3C**). The interface between the PHEV RBD and DPEP1 buries ∼1150 Å^2^ (530 Å^2^ on the RBD and 620 Å^2^ on the receptor), an area considerably smaller than the buried surface area (BSA) of other betacoronavirus RBD/receptor complexes (which are in the range of 1600-2000 Å^2^)^15^. The surface electrostatic potential shows moderate charge complementarity between the PHEV RBD and DPEP1, with most of the RBD interface having no net charge, except for a few positive patches that face negative surfaces on DPEP1 (**Figure 5C**). The complex is stabilized by polar interactions that include five hydrogen bonds, with R130_DPEP1_ and E351_DPEP1_ participating in most of them, and a salt bridge (**Table S2, Figure 5D**). Besides, two aromatic residues from the RBD (F507_RBD_ and W533_RBD_) stack against the long aliphatic side chains of arginine residues from DPEP1 (**Figure 5D**). Altogether, the small BSA of the interface, the limited charge complementarity and the scarcity of polar interactions are consistent with the measured binding affinity in the low micromolar range.

**Figure 5.**
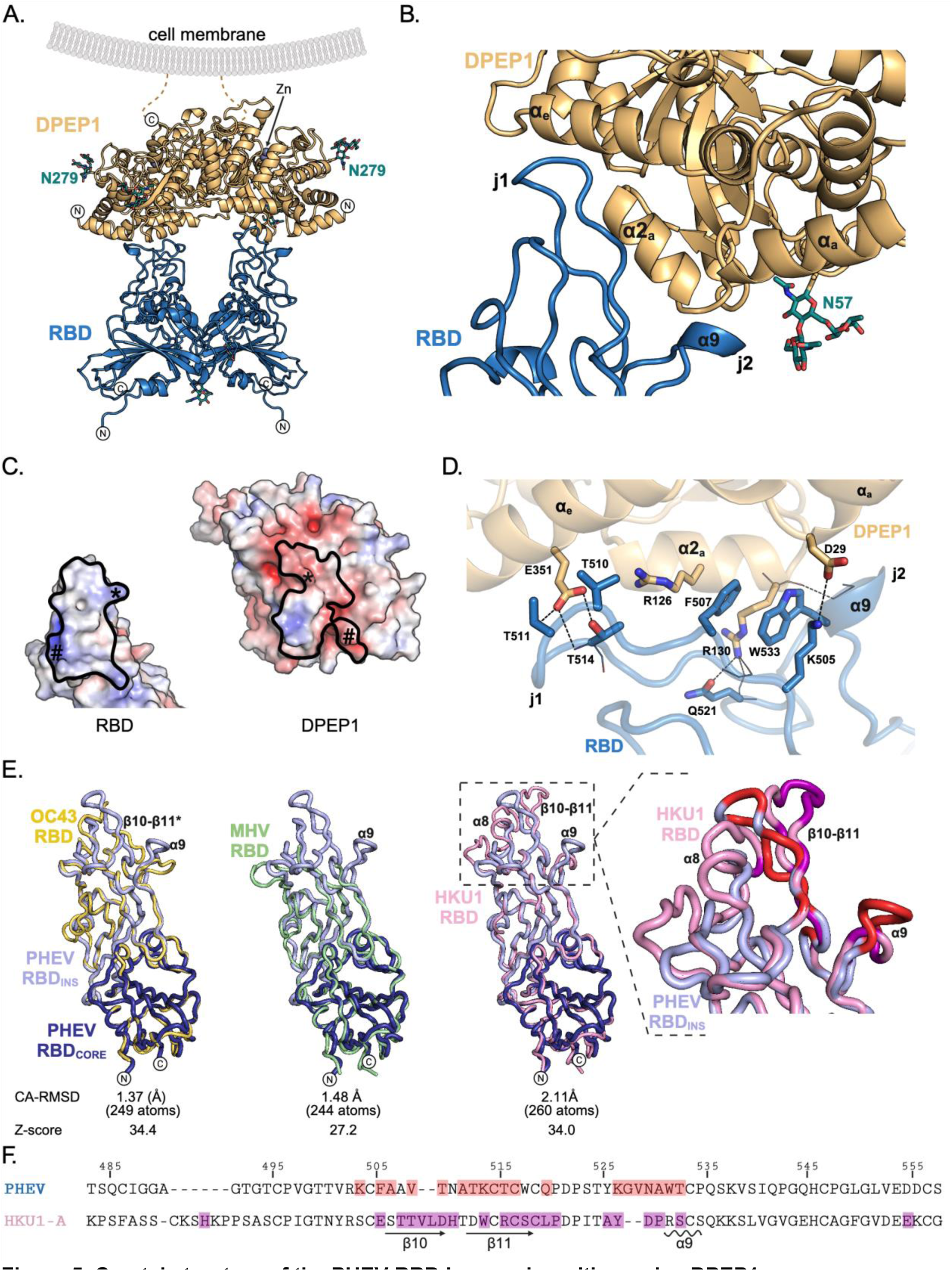
Crystal structure of the PHEV-RBD in complex with porcine DPEP1. **A.** Structure of the isolated RBD/DPEP1 complex. The N- and C-termini of the proteins are indicated with circled ‘N’ and ‘C’ letters, respectively. Glycans are represented with green sticks and the asparagine residues to which they are linked are indicated. The DPEP1 active site contains two zinc ions (gray spheres) and the orientation of the protein with respect to the plasma membrane is shown. The location of the cell membrane on which DPEP1 is inserted is indicated, showing that the RBD recognizes the opposite face. **B.** Close-up view of the RBD-DPEP1 interface. The RBD tip adopts an open mouth shape with two jaws, j1 and j2. Secondary structure elements involved in the interaction are highlighted. The glycan attached to N57 is represented with green sticks. **C.** Open book representation of the PHEV-RBD and DPEP1 showing the surface electrostatic potential. The thick black line delimits the residues involved in the interface. The star (*) and hash (#) signs point out the regions with complementary charge that are in contact. **D.** Detailed atomic interactions at the RBD/DPEP1 interface. The side chains of relevant residues are shown in sticks, while main chain atoms are in thin lines. Dotted lines are used to represent hydrogen bonds or salt bridges. Some secondary structure elements are labeled, as well as jaw 1 (j1) and jaw 2 (j2). **E.** Superposition of the PHEV RBD structure with those from OC43 (yellow, left panel, PDB: 8TZU), MHV (green, central panel, PDB: 6VSJ) and HKU1 (pink, right panel, PDB: 5KWB). The RBD subdomains are shown in the PHEV structure, with the core in dark blue (PHEV RBD_CORE_) and the insertion in light blue (PHEV RBD_INS_). Relevant secondary structure elements are labeled. A star symbol (*) is used to indicate correspondence, even though the secondary structure is different. Below each alignment are the values of the root mean square deviation between alpha-carbons (CA-RMSD) and the Z-score of the pairwise alignment performed with the Dali server (Z>20 indicates that the proteins are structural homologs^59^). The inset on the far-right zooms into the RBD tips (dashed box) from PHEV (light blue) and HKU1 (pink), where the RBD residues buried at the interface formed with DPEP1 and TMPRSS2 are indicated in red and magenta, respectively and relevant secondary structure elements are labeled. **F.** Amino acid sequence alignment of the PHEV and HKU1 RBDs, showing the residues buried by the receptor (shaded in red and violet, respectively). The numbers above the sequence indicate the residue numbering in the PHEV spike. Two arrows and a wavy line below the alignment indicate beta strands and an alpha helix, respectively, in the HKU1 RBD.

### Structural conservation between the RBDs of PHEV and HKU1

Structural alignment of the PHEV RBD with those from OC43^27^ (PDB: 8TZU), MHV^28^ (PDB: 6VSJ) and HKU1^29^ (PDB: 5KWB) shows that all of them share a conserved fold (**Figure 5E**). This is also revealed by low values of root mean square deviation between alpha-carbon atoms (CA-RMSD), and high pairwise alignment Z-scores on the Dali server^30^. This common fold consists of two subdomains: a core (PHEV residues 327-434 and 583-603) that has a topology similar to the RBDs from other betacoronaviruses (SARS-CoV, MERS-CoV) and an insertion (PHEV residues 435-582) that is unique to embecoviruses and encompasses most of the highly divergent region presented above (**Figure S3C**). Several segments of the insertion subdomains from different RBDs superpose well (**Figure 5E**), with the exception of the highly variable region located toward the tip of RBD tip, where the DPEP1 contact residues are located. The RBD tip also harbors the TMPRSS2 binding site in the HKU1 RBD^15^ (**Figure 5E**), and the same structural elements adopt slightly different conformations to participate in the interaction (**Figure 5F**). The RBD sequences from betacoronavirus 1 members have deletions in the positions corresponding to the receptor-binding site (**Figure S5**), which could be responsible for their different receptor specificities. The most extensive deletion is found on the OC43 variable region, which lacks the β10-β11 hairpin and the segment corresponding to α8 protrudes on top of the resulting structure (**Figure 5E, Figure S5**).

### Functional identification of critical residues for DPEP1-PHEV spike interaction

To functionally test the role of specific residues at the PHEV spike/DPEP1 interface in virus entry, we performed site-directed mutagenesis of the PHEV spike and porcine DPEP1 and evaluated the ability of these mutants to mediate pseudotype entry and cell-cell fusion (**Figure 6A-B**). First, we mutated ten interface residues of the PHEV spike and included variant Q487A (a residue located in the loop below j1 and not in contact with DPEP1) as control. All spike mutants were incorporated correctly into pseudotypes (**Figure 6A**, left). Strikingly, the F507R and W533A spike mutants almost completely abolished PHEV pseudotype infectivity in porcine DPEP1-expressing cells (**Figure 6A**, middle) and led to a >10-fold reduction in cell-cell fusogenicity (**Figure 6A**, right). Other mutations (V510R, T511D, T514A and T517R) decreased pseudotype infectivity to a lower extent (5- to 38-fold) without decreasing cell-cell fusion. We then performed similar analyses by mutating three interface residues of DPEP1 (T123, R126 and E351). We also used mutant Q42A, a residue away from the RBD recognition site, as control. All DPEP1 mutants were well-expressed in HEK293T cells (**Figure 6B**, top). The E351R mutation completely abolished entry of a PHEV pseudotype (**Figure 6B**, bottom left) and strongly reduced cell-cell fusion (**Figure 6B**, bottom right). Mutating this residue to alanine (E351A) also slightly decreased PHEV pseudotype entry but did not alter cell-cell fusogenicity. This residue is therefore critical for the PHEV spike-DPEP1 interaction.

**Figure 6.**
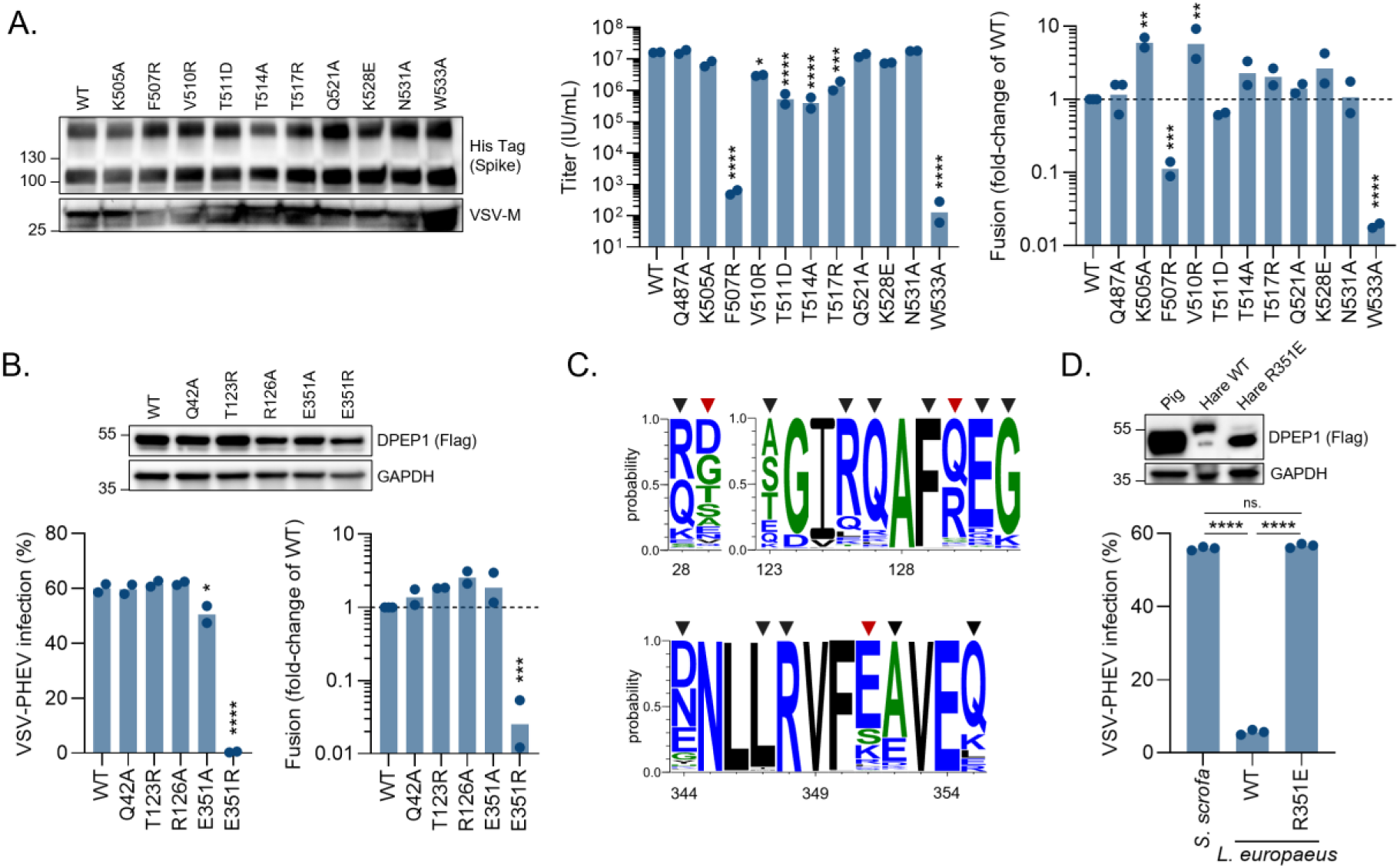
Identification of critical residues for the interaction between porcine DPEP1 and the PHEV spike. **A.** Left: western blot confirmation of the incorporation of the WT and mutant PHEV spikes into VSV pseudotypes. A His-Tag added to the spike C-terminus was used for detection and VSV-M was used as a loading control. Middle: PHEV pseudotypes were produced with WT or the indicated mutant spikes and titrated on HEK293T cells expressing DPEP1. Pseudotype titers are shown. Right: HEK293T GFP-Split cells were transfected with porcine DPEP1 and WT or mutant PHEV spikes. The fold-change in cell-cell fusion relative to the WT is shown. **B.** Top: western blot confirmation of WT and mutant DPEP1 expression. A Flag-Tag added to the DPEP1 C-terminus was used for detection and GAPDH was used as a loading control. Bottom left: HEK293T were transfected with wild-type (WT) or mutant DPEP1 and infected with PHEV pseudotypes. Percentages of infection are shown. Bottom right: HEK293T GFP-Split cells were transfected with the PHEV spike and WT or mutant DPEP1. The fold-change in cell-cell fusion relative to the WT is shown. **C.** Probability of different amino acids occupying the indicated positions (numbers at the bottom) of mammalian DPEP1 orthologs (n = 225 mammal species). Inverted triangles indicate the residues involved in the interaction with the PHEV spike, with those in red forming polar interactions. **D.** HEK293T cells were transfected with the pig (*S. scrofa*, E351) or hare (*L. europaeus*), either WT (R351) or mutant (R351E), and infected with PHEV pseudotypes. The percentage of infected cells is shown. The statistical significance of two-sided unpaired t-tests are indicated: **** P < 0.0001, ns: not significant. The bar represents the mean and each dot is a technical replicate (n = 3). In **A** and **B**, the bar represents the mean, each dot represents an independent experiment (n = 2) and the statistical significance of a one-way ANOVA with Dunnett’s multiple comparison test on log-transformed data is shown: * P < 0.05, ** P < 0.01, *** P < 0.001, **** P < 0.0001.

### Cross-species variability in critical DPEP1 residues may explain PHEV susceptibility

We then investigated whether interspecies variations in contact residues (**Figure S4**) could alter the ability of diverse mammalian DPEP1 orthologues to allow PHEV entry. We first examined the degree of conservation of DPEP1 interface residues in 225 mammalian species (**Figure 6C**). Some contact residues were strongly conserved (e.g. F129, L347, R348), but the majority, including the three residues involved in polar contacts with the PHEV RBD (D29, R130, E351), showed some degree of variability (**Figure S4**). In particular, some species such as the European hare (*Lepus europaeus*) have an arginine at position 351 (R351). In agreement with our mutagenesis results, the hare DPEP1 orthologue (R351) was poorly able to mediate entry of PHEV pseudotype into HEK293T cells (**Figure 6D**). Mutating the arginine to a glutamic acid (R351E) restored the ability of the hare DPEP1 to mediate PHEV entry to the same level as porcine DPEP1 (**Figure 6D**). These results confirm the functional importance of this residue for the interaction with the PHEV spike and suggest that it could be a determinant of PHEV cross-species transmissibility.

### PHEV spike opening does not depend on sialoglycan binding

Various studies have reported that in the absence of sialic acid ligand, the embecovirus spike protein (from MHV, HKU1, OC43) adopts a closed conformation^4–6^. Binding of the HKU1A spike NTD to specific sialoglycan ligands triggers an allosteric conformational change that promotes RBD opening^7^ and exposure of the protein-receptor binding site^15^. To explore whether this is the case for PHEV, we studied its spike ectodomain by cryo-EM. The construct used was stabilized by mutation of the furin site (^751^RSRR^754^ to ^751^GSAG^754^) to impair cleavage, and by fusing a “foldon” trimerization motif^31^ at the C-terminus, without the stabilizing proline mutations in the spike S2 subunit commonly used for structural studies of the spike^32,33^. Analysis of the cryo-EM data (**Figure S6**) indicated that the PHEV spike can display the RBD in different configurations, with 16% of the spikes in the closed form (three RBDs in the down position) (**Figure 7A**). Upon non-uniform refinement of this class, we obtained a cryo-EM map at 3.4 Å resolution (**Table S3**), which allowed us to build an atomic model of the closed spike (**Figure 7B**). Despite the good overall resolution, the local density of the NTD upper subdomain (which contains the glycan binding site) and the RBD tip (at the location of the insertion subdomain) were weak (**Figure S6**), precluding model building in these regions. Furthermore, most of the analyzed particles belonged to conformations where the spike had 1-RBD (26%), 2-RBDs (43%) or even 3-RBDs (15%) in the ‘up’ position (**Figure 7A**, **Figure S6**, **Table S3**). As the number of up-RBDs increases, their density becomes weaker (identified in **Figure 7A** as ‘up/missing’), indicating that they are highly flexible in this conformation. Importantly, the identification of partially and fully open conformations shows that the PHEV spike does not require glycan binding to trigger opening, in contrast to the HKU1A spike. This interpretation is consistent with our BLI experiments showing that the spike ectodomain readily binds to DPEP1 (**Figure 3F**).

**Figure 7.**
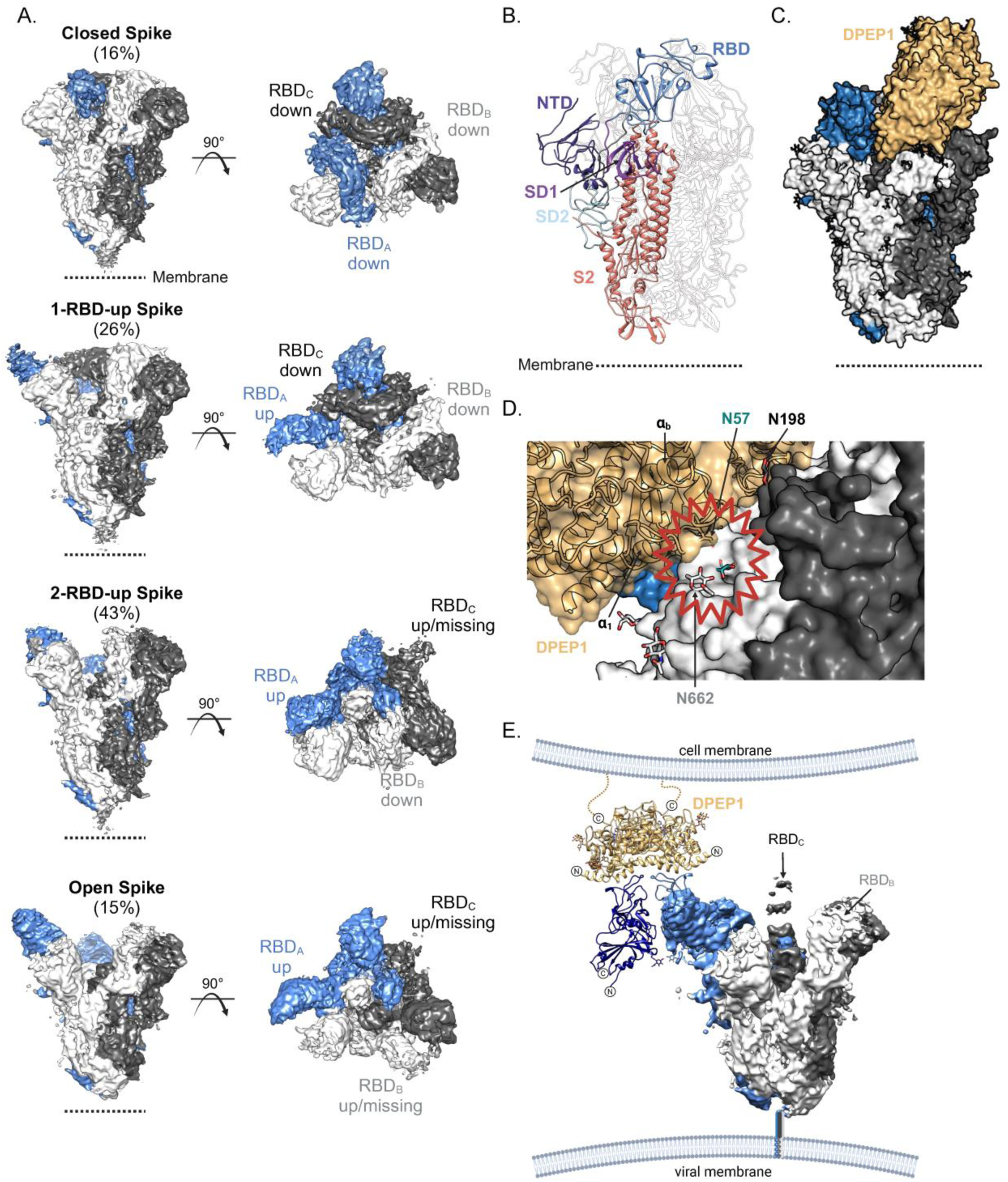
Cryo-electron microscopy (cryo-EM) of the PHEV spike ectodomain. **A.** Refined cryo-EM maps of different spike conformations obtained after 3D classification (**Figure S6**). The left panels show side views of the spike where each protomer is labeled with a subscript (A, B, C) and is shown in a different color (blue, white and dark gray). A dashed line in each panel indicates the position of the viral membrane. The right panels show a top view of the spike trimers. Each conformation is identified with the number of RBDs in the ‘up’ position and the number between parenthesis indicates the percentage of particles in each state. **B.** Cartoon representation of the atomic model of the PHEV spike ectodomain in the closed state. For clarity, two protomers are outlined and the third one shows the protein domains in different colors. **C.** Surface representation of the PHEV spike with porcine DPEP1 upon superposition of the RBD-DPEP1 crystal structure on the closed RBD_A_ (blue). **D.** Zoom into the contact region between porcine DPEP1 and the closed spike upon superposition of the RBD-DPEP1 crystal structure on RBD_A_. DPEP1 is shown with transparent surface to facilitate the identification of the indicated secondary structure elements. Glycans from DPEP1 are shown in green, with the one attached to N57 penetrating the spike. Glycans from the spike protomer B are indicated in white sticks, while those in dark gray are on protomer C. The region where DPEP1 clashes with the spike is highlighted with a wavy red outline. **E.** Fitting of the RBD/DPEP1 crystal structure into the density map of the open spike. N- and C-termini are indicated with circled ‘N’ and ‘C’ letters, respectively. DPEP1 is shown with the two RBD molecules found in the asymmetric unit. The viral and cell membranes are drawn to indicate the topology of the complex.

### DPEP1 binds only open spikes

Finally, to understand whether spike opening is required for DPEP1 binding, we superimposed the RBD/DPEP1 crystal structure onto the closed spike (**Figure 7C**) and evaluated the potential interaction of the receptor with down-RBDs on the trimer. This exercise showed that DPEP1 cannot bind to a closed spike (for example, to RBD_A_) because its α_b_-α_1_ loop and the glycan attached to N57 in DPEP1 would clash with the counterclockwise down RBD from a neighboring protomer (RBD_B_; **Figure 7D and S4**). Further steric hindrance could be generated by the spike glycans at N198 and N662, which would lie below DPEP1 (**Figure 7D**). We therefore conclude that DPEP1 can bind only to RBDs in the ‘up’ position and that interaction with the receptor stabilizes open forms of the spike. Moreover, fitting the crystal structure of the RBD/DPEP1 complex into the cryo-EM density map of the open spikes further shows that the two protomers of the DPEP1 dimer cannot be reached by the RBDs of a single spike (**Figure 7E**), indicating potential linkage of the spikes by the dimeric receptor, as recently shown for SARS-CoV- _234._

## Discussion

PHEV circulates subclinically in most swine herds worldwide but causes gastrointestinal and neurologic symptoms in immunologically naive animals, being typically fatal in piglets less than 3-4 weeks of age^35^. Therefore, in the absence of a vaccine or treatment, PHEV poses an economic threat to swine herds. The identification of DPEP1 as a functional PHEV receptor may facilitate the development of new antivirals targeting entry. Previous work showed the involvement of attachment factors in viral entry, including 9-*O*-acetylated sialic acids, heparan sulfates, the neural cell adhesion molecule (NCAM), and the MERS-CoV receptor DPP4^8,36–38^, but *bona fide* proteinaceous receptors were unknown for PHEV and other members of the betacoronavirus 1 species.

In humans, the main function of DPEP1 is to hydrolyze several dipeptides (e.g. glutathione) in renal metabolism. To our knowledge, DPEP1 has not been involved previously in viral infections. In pigs, DPEP1 is mainly expressed in the kidney and gastrointestinal tract^39^, which is consistent with the gastrointestinal symptomatology of PHEV infection. Although PHEV is also neurotropic, DPEP1 does not appear to be highly expressed in the pig brain^39^. Whether suboptimal DPEP1 expression in the brain is sufficient to allow PHEV neurotropism - or whether PHEV can use a different receptor in brain cells - deserves further investigation. Future *in vivo* work will help to unravel the role of DPEP1 in PHEV replication and pathogenesis.

Although viruses from the same species usually use the same receptor for viral entry, our study showed that the PHEV spike is the only betacoronavirus 1 spike tested that interacts with DPEP1. Diversity in receptor usage among members of the betacoronavirus 1 species is likely related to the complex evolutionary history of coronaviruses, which combines a high mutation rate, extensive recombination, frequent host jumps, and host adaptation. Importantly, how DPEP1 usage was acquired by PHEV during evolution of the betacoronavirus 1 species remains an open question. One possibility is that PHEV acquired its RBD by recombination with another virus, potentially using DPEP1 itself. We were unable to identify a clear candidate for the origin of the PHEV RBD. Therefore, more work, including novel virus discovery, is needed to understand how and when PHEV acquired the ability to use DPEP1 as a receptor and whether it can be used by other viruses.

Although the various members of the betacoronavirus 1 species are host-specific, we found that the PHEV spike is compatible with the human orthologue of DPEP1 and other entry determinants, since PHEV pseudotypes infected one third of the human cell lines tested. In contrast, other DPEP1 orthologues did not allow PHEV entry (e.g. European hare). The extent of cross-species compatibility of the PHEV spike with mammalian DPEP1 proteins needs to be further explored, as this would help to assess the cross-species transmission potential of PHEV. Indeed, despite widespread subclinical circulation of PHEV in pig farms, its ability to infect other mammals, including humans, remains unclear.

Our results strongly argue against a conserved entry mechanism for all embecoviruses. For instance, despite the preservation of the HE-coding gene, and in clear contrast with HKU1, OC43 and BCoV, PHEV does not require sialic acid for entry. Previous studies have shown that the sialoglycan binding site is in the spike NTD and that all interacting residues are conserved OC43, BCoV, HKU1 and PHEV^5^. However, the PHEV spike was shown to have a 1,450- and 45-fold lower affinity for sialoglycans compared to those of BCoV and OC43, respectively^8^, suggesting that glycans may play a less prominent role in regulating PHEV entry. The W89A mutation in the NTD makes the HKU1 spike unable to bind sialoglycans^8^, adopt open conformations^7^, and mediate fusion^15^. Similarly, the W90A mutant BCoV pseudotypes are non-infectious^8^. Although the equivalent W90A mutation in the PHEV spike abrogates interaction with glycans^8^, we show here that it does not prevent infection of target cells. Moreover, neuraminidase treatment of target cells did not abrogate viral entry. Our cryo-EM analyses also showed that the PHEV spike trimer samples a conformational landscape of closed, partially and fully open forms in the absence of ligands, which has not been observed in spike proteins from other embecoviruses. The independence of glycan binding, with the concomitant high frequency of RBD opening, may be a mechanism to compensate for the low affinity of the PHEV RBD for DPEP1.

Our work supports that, instead of the two-step entry mechanism proposed for HKU1, PHEV follows a model similar to that of SARS-CoV-2, depending exclusively on a protein receptor that binds to spontaneously “up” RBDs. Nevertheless, our work reveals a conserved binding mode among RBDs from embecoviruses, despite the observed diversity in RBD sequences and receptor specificity. The structural organization of the RBD confers it a remarkable versatility in receptor binding, whereby completely different sequence motifs can be accommodated into the tip, enabling changes in receptor usage. In particular, the deletions present in the sequences of the betacoronavirus 1 RBD tips may be responsible for the receptor usage diversity within this species.

In sum, we have identified a new coronavirus receptor and shown that embecoviruses have divergent mechanisms to regulate the opening of the spike protein, yet they use the tip of the RBD to recognize different receptors. This parallels the ability of other betacoronaviruses like sarbeco- and merbecoviruses to adapt a common RBD scaffold to different receptors. Embecoviruses seem to have achieved such versatility by exchanging only small pieces that are confined to the RBD tip, as opposed to the larger variable region observed in other betacoronaviruses.

Finally, this study expands the repertoire of membrane-bound peptidases that function as coronavirus receptors. The reason why coronaviruses preferentially use peptidases for entry remains elusive, as their catalytic activity was shown not to no be required, but it seems unlikely that it is only a coincidence.

## Methods

### Cell lines

The NCI-60 panel (dtp.cancer.gov/discoverydevelopment/nci-60) was obtained from the National Cancer Institute. The panel consists of 60 well-characterized cancer cell lines from various origins. 48 adherent cell lines were included in our analysis. Their identity was confirmed by Short Tandem Repeat (STR) genotyping, as previously described^18^. NCI-60 cell lines were cultured in RPMI (Gibco) supplemented with 10% FBS (Gibco), 10 units/mL penicillin, 10 µg/mL streptomycin (Gibco), 250 ng/mL amphotericin B (Gibco) and 5 µg/mL prophylactic plasmocin (InvivoGen). HEK293T-GFP1-10 and HEK293T-GFP11 cells were kindly provided by Olivier Schwartz (Institut Pasteur, Paris, France) and were cultured in DMEM supplemented with 10% fetal bovine serum (FBS), 1% non-essential amino acids, 10 U/mL penicillin, 10 µg/mL streptomycin, 250 ng/mL amphotericin B, and 1 µg/mL of puromycin (Gibco). PK-15 (Cytion) and swine primary kidney cells were cultured in MEM (Gibco) supplemented with 10% fetal bovine serum (FBS), 10 U/mL penicillin, 10 µg/mL streptomycin, and 250 ng/mL amphotericin B. All cell lines were grown at 37°C and 5% CO_2_ and were regularly shown to be free of mycoplasma contamination by PCR.

### Plasmids

The genes encoding the full PHEV (GenBank DQ011855.1), OC43 (RefSeq NC_006213.1), BCoV (RefSeq NC_003045.1), ECoV (GenBank EF446615.1), CRCoV (GenBank EU983106.1) and HKU24 (RefSeq NC_026011.1) spikes were codon-optimized for human expression, synthesized, and cloned into a pcDNA3.1 vector bearing a C-terminal 6xHis tag (Genscript). To obtain the human DPEP1 and TMPRSS2 coding sequences, total RNA was extracted from HEK293T or SW-620 cells, respectively, reverse-transcribed using Oligo(dT) and Superscript IV (Invitrogen), amplified by PCR using Phusion Hot Start II DNA polymerase (Thermo Scientific™) with specific primers (DPEP1: 5’-ttaaacttaagcttgccaccatgtggagcggatggtggctg-3’; 5’-tcgtcgtcatccttgtaatccaggagagacagacagaggacc-3’; TMPRSS2: 5’-ttaaacttaagcttgccaccatggctttgaactcaggg-3’, 5’-tcgtcgtcatccttgtaatcgccgtctgccctcatttg-3’), and cloned into a pcDNA3.1-Flag vector using the NEBuilder^®^ HiFi DNA Assembly kit (New England Biolabs) following the manufacturer’s instructions. The pig (*Sus scrofa*, RefSeq NM_214108.1) and hare (*Lepus europaeus*, RefSeq XM_062176926.1) orthologs of DPEP1 were synthesized and cloned into a pcDNA3.1-Flag vector (Genscript). DPEP1 and PHEV spike mutants were obtained by site-directed mutagenesis using the QuikChange II XL Site-Directed Mutagenesis Kit (Agilent) according to the manufacturer’s instructions. Mutagenesis primers are listed in **Table S4**. Successful mutagenesis was confirmed by whole plasmid sequencing (Plasmidsaurus or Eurofins).

### Phylogenetic analysis of embecovirus spikes

Multiple sequence alignments of embecovirus spike proteins were performed using the Clustal omega tool in Molecular Evolutionary Genetics Analysis Version 11 (MEGA11) software with default parameters^54^. The evolutionary history was inferred by using the Maximum Likelihood method and JTT matrix-based model^55^. The tree with the highest log likelihood is shown. Initial tree(s) for the heuristic search were obtained automatically by applying Neighbor-Join and BioNJ algorithms to a matrix of pairwise distances estimated using the JTT model, and then selecting the topology with superior log likelihood value. The tree was drawn to scale, with branch lengths measured in the number of substitutions per site. Bootstrap analysis was used to test the robustness of the tree topology (100 resamplings). Spike protein similarity plots were built using the SimPlot++ software^40^, with a 100 amino acid sliding window and a 10 amino acid step.

### VSV pseudotyping

T75 flasks were coated with poly-D-lysine (Gibco) for 2 h at 37°C, washed with water, and seeded with 8 × 10^6^ HEK293T cells. The following day, cells were transfected with 30 µg of plasmid encoding for PHEV S, VSV-G or an empty vector using Lipofectamine 2000 (Invitrogen) following the manufacturer’s instructions. At 24 h post-transfection, cells were inoculated at a multiplicity of infection (MOI) of 3 infectious units per cell for 1 h at 37°C with a VSV encoding GFP, lacking the glycoprotein gene G (VSVΔG-GFP), and previously pseudotyped with G. Cells were washed three times with PBS and 8 mL of DMEM supplemented with 2% FBS were added. Supernatants containing pseudotypes were harvested 24 h later, cleared by centrifugation at 2,000 g for 10 min, passed through a 0.45 µm filter, aliquoted, and stored at -80°C.

### Infection of the NCI-60 panel

Cells were seeded in 96-well plates the day before the infection. Pseudotypes were mixed 1:1 with an anti-VSV-G monoclonal antibody to remove residual VSV-G and incubated for 20 min at 37°C. Cell culture medium was removed and cells were inoculated with 50 µL of the antibody-treated pseudotypes. Plates were incubated for 2 h at 37°C and 50 µL of RPMI supplemented with 5% FBS were added to each well. After 20 h, cells were imaged in the Incucyte SX5 Live-Cell Analysis System (Sartorius). The infected cell percentage was calculated as the ratio between cell confluence and the percentage of GFP-positive area, both quantified automatically with the Incucyte Analysis software. Cells were also infected with a bald pseudotype to measure the background signal resulting from cell auto-fluorescence or residual VSV-G-pseudotyped particles. The values obtained in these negative controls were subtracted from the corresponding PHEV pseudotype measurements. All corrected infection values below 0.1% were set to 0.1% (infectivity threshold).

### Gene correlation analyses

We normalized infectivity data as previously described^18^. The proportion of infected cells, Q, was measured as the ratio between the GFP area and cell confluence. We transformed Q-values into multiplicities of infection (MOIs) as follows: MOI = –ln(1-Q). Relative MOI values were then calculated as R = 100 × MOI/max(MOI). Finally, values were log-transformed as log_2_(R+1), and the two log_2_(R+1) data obtained in each experimental replicate were averaged. Processed RNA-seq and microarray datasets were downloaded from the CellMiner website (discover.nci.nih.gov/cellminer/loadDownload.do, “RNA-seq - composite expression” and “Agilent mRNA - log2” files). These transcriptomic data were available for all cell lines except MDA-MB-468. RNA-sequencing data were available as log_2_(rpkm+1) and microarray data as log_2_(intensity values). For each gene, RNA-seq and microarray data were first separately scaled as a percentage of the maximal expression value observed among the 47 cell lines. For each cell line-gene combination, scaled RNA-seq and microarray data were then averaged. The average expression data of each gene encoding a membrane-associated protein (7694 genes) was correlated to log_2_(R+1) infectivity data across the 47 cell lines using a Pearson correlation. Bonferroni multiple test correction was applied to identify genes whose expression correlated significantly with infection levels.

### Cell-cell fusion assay

The cell-cell fusion assay was performed as previously described^41^. Briefly, HEK293T-GFP1-10 and HEK293T-GFP-11 were mixed at a 1:1 ratio (total of 6 × 10^4^ cells per well of a 96-well plate) and were co-transfected with PHEV spike- and DPEP-expressing plasmids (or empty vectors) at a 1:1 ratio (100 ng total DNA) using Lipofectamine 2000 (Invitrogen) following the manufacturer’s instructions. For the TMPRSS2 assays, cells were transfected with spike-, DPEP1-, and TMPRSS2-expressing plasmids (or empty vectors) at a 1:1:1 ratio (100 ng of total DNA). Plates were placed at 37°C and 5% CO2 in an Incucyte SX5 LiveCell Analysis System (Sartorious). GFP signal and phase contrast images were analysed at 18-24 h post-transfection. The percentage of fusion was calculated as the ratio between GFP area and cell confluence, both quantified automatically with the Incucyte Analysis software.

### Neuraminidase treatment

HCT-15 cells were plated in a 96-well plate in presence or absence of 40 µg/mL of neuraminidase (NA) from *Clostridium perfringens* (Sigma-Aldrich). The next day, cells were infected with PHEV or VSV-G pseudotypes as described above, in the presence or absence of 40 µg/mL NA. Cells were also infected with a GFP-expressing IAV (strain PR8) as a positive control for sialic acids depletion. After 18-24 h, cells were imaged in the Incucyte SX5 Live-Cell Analysis System (Sartorius). The infected cell percentage was calculated as the ratio between cell confluence and the percentage of GFP-positive area, both quantified automatically with the Incucyte Analysis software.

### Pseudotype infectivity competition assay

PHEV pseudotypes were pre-incubated 1:1 with anti-VSV-G antibody and with serial dilutions of soluble human DPEP1 (Sino Biologicals) or vehicle (H_2_O) for 1 h at 37°C, and then used to infect either HCT-15 or DPEP1-transfected HEK293T cells seeded the previous day in 96-well plates. The percentage of infected cells at 18-24 hpi was measured using the Incucyte SX5 Live-Cell Analysis System (Sartorious) and the Incucyte analysis software.

### Flow cytometry measurement of DPEP1 binding

HEK293T cells were seeded in 6-well plates. The next day, they were transfected with 2.5 µg of spike-encoding plasmid using Lipofectamine 2000 following the manufacturer’s instructions. After 24 h, cells were detached gently with cold staining buffer (PBS, 0.5% BSA, 2mM EDTA) and incubated with serial dilutions of soluble recombinant human DPEP1 (Sino Biologicals) for 30 min at 4°C. Cells were washed with PBS and incubated with an anti-DPEP1 antibody (Atlas Antibodies, HPA012783, dilution 1:100) for 30 min at 4°C. After another wash with PBS, cells were incubated with a goat anti-rabbit IgG-AF647 secondary antibody (Invitrogen, A32733, dilution 1:400) for 30 min at 4°C. Cells were fixed with 4% paraformaldehyde (PFA) for 10 min at room temperature, washed with PBS and resuspended in staining buffer. Cells were analyzed on a FACSVerse™ (BD Biosciences) flow cytometer and results were analyzed using the FlowJo software (v10.10). The median fluorescence intensity (MFI) obtained at each soluble DPEP1 concentration was normalized to the one obtained in the absence of soluble DPEP1.

### Full virus competition assay

PHEV (strain 67N) was obtained from the National Veterinary Services Laboratories (NVSL; United States Department of Agriculture [USDA], Ames, IA, USA) and was propagated in swine kidney primary cells (SPKC) as previously described^52^. For the competition assay, SPKC and PK-15 cells were plated in a 12-well plate. The next day, the virus was incubated with soluble DPEP1 (Sino Biologicals) or vehicle (H_2_O) at a concentration of 25 µg/mL for 30 min at 37°C. Cells were then inoculated with 100 µL of treated virus and incubated for 2 h at 37°C. Cells were washed three times with PBS and 1 mL of MEM supplemented with 2% FBS was added. Supernatants were harvested at 2, 12, 24, and 48 h post-inoculation and viral RNA was extracted using the NZY Viral RNA Isolation kit (NZYtech) according to the manufacturer’s instructions. Viral RNA (PHEV N) was quantified by RT-qPCR using previously published primers and probe^42^, and the TaqPath™ 1-Step RT-qPCR Master Mix (Applied Biosystems™) according to the manufacturer’s instructions.

### Western blot

A 1 mL volume of supernatant containing pseudotype was pelleted by centrifugation at 30,000 g for 2 h at 4°C and lysed in 30 µL of NP-40 lysis buffer (Invitrogen) for 30 min on ice. Around 5 × 10^5^ transfected cells were lysed in 50 µL of NP-40 lysis buffer (Invitrogen) for 30 min on ice. Lysates were cleared by centrifugation at 15,000 g for 10 min at 4°C. Cleared lysates were mixed with 4X Laemlli buffer (Bio-Rad) supplemented with 10% β-mercaptoethanol and denatured at 95°C for 5 min. Proteins were separated by SDS-PAGE on a 4-20% Mini-PROTEAN TGX Gel (Bio-Rad) and transferred onto a 0.45 µm PVDF membrane (Thermo Scientific). Membranes were blocked for 1 h at room temperature in TBS-T (20 mM tris, 150 mM NaCl, 0.1% Tween-20, pH 7.5) supplemented with 3% Bovine Serum Albumin (BSA; Sigma). Membranes were then incubated for 1 h at room temperature with the following primary antibodies: mouse anti-His-Tag (dilution 1:1,000, clone HIS.H8, Invitrogen MA121315), mouse anti-Flag (dilution 1:1000, clone M2, Sigma-Aldrich F1804), mouse anti-VSV-M (dilution 1:1000, clone 23H12, Kerafast EB0011), rabbit anti-GAPDH (dilution 1:3,000, Sigma-Aldrich ABS16). Membranes were washed 3 times with TBS-T and incubated for 1 h at room temperature with an HRP-conjugated anti-mouse (dilution 1:50,000, Invitrogen, G-21040) or anti-rabbit (dilution 1:50,000, Invitrogen, G-21234) secondary antibody. After 3 washes in TBS-T, the signal was revealed with SuperSignal West Pico PLUS (Thermo Scientific) following the manufacturer’s instructions. Images were acquired on an ImageQuant LAS 500 (GE Healthcare) and analyzed with Fiji software.

### Construct design for protein expression and purification

Codon-optimized synthetic genes coding for the PHEV spike (residues 15-1274, NCBI accession AAY68297.1), porcine DPEP1 (residues 17-385, GenBank accession CAA37762.1, UniProt P22412) and SARS-CoV-2 Omicron BA.4/5 spike^43^ (residues 1-1208, Wuhan numbering) were purchased to Genscript. Cloning of these genes was also performed by Genscript. Plasmids coding for the SARS-CoV-2 Wuhan RBD and soluble ACE2 have been previously described^14^.

The PHEV-spike ectodomain was stabilized in the prefusion form by introducing mutations in the furin site (^751^RSRR^754^ to ^751^GSAG^754^) and adding a Foldon (YIPEAPRDGQAYVRKDGEWVLLSTFL) trimerization motif at the C-terminus. This construct was cloned into pCAGGS with an Ig kappa signal peptide (METDTLLLWVLLLWVPGSTG) and a thrombin cleavage site at the C-terminus (LVPRGS) followed by a His-(HHHHHHHH), a Strep-(WSHPQFEK) and an Avi-tag (GLNDIFEAQKIEWHE). This plasmid was used as a template to amplify the PHEV-RBD (residues 327-605) coding sequence, which was cloned into pCAGGs with the same signal peptide and tags as the spike.

The SARS-CoV-2 Omicron BA.4/5 spike ectodomain was stabilized with six proline mutations in S2 (equivalent to F817P, A892P, A899P, A942P, K986P and V987P, Wuhan numbering), the abrogation of furin cleavage (^682^RRAR^685^ to ^682^GSAS^685^, Wuhan numbering) and the addition of a C-terminal Foldon motif. This construct was cloned in the pCI-Neo plasmid, followed by His, Strep and Avi tags.

Soluble DPEP1 was cloned into a modified pcDNA3.1(+) vector, which contains a CMV exon-intron-exon sequence to boost expression, downstream of the CD5 signal peptide (MPMGSLQPLATLYLLGMLVASCLG) and upstream of an enterokinase cleavage site (DDDDK) and a double Strep tag. Another DPEP1 construct (DPEP1-Avi) was designed with a single Strep sequence followed by the Avi tag after the enterokinase site.

For expression in insect cells, the PHEV-RBD and DPEP1 were cloned into a modified pMT/BiP plasmid (Invitrogen; hereafter termed pT350), which translates the protein in frame with an enterokinase cleavage site and a double strep-tag at the C-terminal end.

### Protein expression and purification

#### Protein expression and purification for X-ray crystallography

Plasmids encoding the PHEV-RBD or DPEP1 were co-transfected with the pCoPuro plasmid for puromycin selection in Drosophila Schneider line 2 cells (S2) using the Effectene transfection reagent (Qiagen). The cell lines underwent selection in serum-free insect cell medium (HyClone, Cytiva) containing 7 μg/ml puromycin and 1% penicillin/streptomycin. For protein production, the cells were grown in spinner flasks until the density reached 10^7^ cells/mL, at which point the protein expression was induced with 4 μM CdCl_2_. After 6 days, the cultures were centrifuged, and the supernatants were concentrated and used for affinity purification in a Strep-Tactin column (IBA). The strep tags were removed by incubating the proteins with 48-60 units of Enterokinase light chain (BioLabs) in the elution buffer supplemented with 2 mM CaCl_2_, at room temperature, overnight. The proteolysis reactions were buffer-exchanged into 10 mM Tris, 100 mM NaCl, pH 8.0, and subjected to a second affinity purification, recovering the flow-through fraction containing the untagged proteins. The proteins were concentrated and the enzymatic deglycosylation with endoglycosidase D (EndoD, 500 units) and endoglycosidase H (EndoH, 1000 units) was set up at room-temperature following overnight incubation in 50 mM Na-acetate, 200 mM NaCl, pH 5.0. The proteins were further purified on a size exclusion chromatography (SEC) Superdex 200 16/600 (Cytiva) column in 10 mM Tris, 100 mM NaCl, pH 8.0, and concentrated in VivaSpin concentrators. The purity of the final protein samples was analyzed by SDS-PAGE followed by Coomasie Blue staining.

#### Purification of complexes used for crystallization screenings

The PHEV-RBD was incubated with DPEP1 (after EK cleavage and deglycosylation) at final concentrations of 150 μM and 42.3 μM (dimer), respectively. Following over-night incubation at 4°C, the reaction was loaded onto a Superdex 200 10/300 increase column (Cytiva) equilibrated in 10 mM Tris-HCl, 100 mM NaCl (pH 8.0) to isolate the complex by SEC. Eluted fractions were analyzed by SDS-PAGE and those corresponding to the ternary complex were pooled, concentrated to 8.8 mg/mL and used in crystallization trials.

### Protein expression and purification for biophysical assays and cryo-electron microscopy

DPEP1, DPEP1-Avi, ACE2, PHEV-RBD, SARS-CoV-2 RBD, PHEV-spike and SARS-CoV-2 spike ectodomain-encoding plasmids were transiently transfected into Expi293F^TM^ cells (Thermo-Fischer) using FectoPro^®^ DNA transfection reagent (PolyPlus). After 5 days at 37 °C, cells were harvested by centrifugation and proteins from the supernatants were purified by affinity chromatography in a Strep-Tactin column (IBA). The eluted fractions were pooled, concentrated and loaded onto a Superdex 200 10/300 increase column (Cytiva) (RBD, DPEP1) or a Superose 6 10/300 increase column (Cytiva) (spike) that had been previously equilibrated in 10 mM Tris-HCl, 100 mM NaCl (pH 8.0). Fractions from each main peak were concentrated and frozen. The purity of the final protein samples was analyzed by SDS-PAGE followed by Coomasie Blue staining.

### Crystallization and structural determination

Crystallization screening trials were carried out by the vapor diffusion method using a Mosquito TM nanodispensing system (STPLabtech, Melbourn, UK) following established protocols^44^. The best crystals of the PHEV-RBD/DPEP1 complex were obtained in 0.1 M 2-(N-morpholino)ethanesulfonic acid (MES, pH 6.0), 0.2 M zinc acetate, 10% w/v polyethylene glycol (PEG) 8000 at 18°C using the sitting-drop vapor diffusion method. Crystals were flash-frozen by immersion into a cryo-protectant containing the crystallization solution supplemented with 33% (v/v) ethylene glycol (EG), followed by rapid transfer into liquid nitrogen. The X-ray diffraction data were collected at 100 K at the Proxima-2A beamline of the SOLEIL synchrotron source (Saint Aubin, France)^45^. Data were processed, scaled and reduced with XDS^46^ and AIMLESS^47^. The structures were determined by molecular replacement using Phaser from the PHENIX suite^48^ with search ensembles obtained from AlphaFold3^25^ (PHEV-RBD) and the crystal structure of human DPEP1 (PDB: 1ITQ). The final models were built by combining real space model building in Coot^49^ with reciprocal space refinement with phenix.refine. The final model was validated with Molprobity^50^. The analyses of the macromolecular surfaces were carried out in PDBePISA^51^. Figures were created using Pymol^52^, Chimera^53^ and BioRender.com.

### Sample preparation for cryo-EM

3 μL of the PHEV-spike ectodomain (0.6 µM trimer) were added to Quantifoil R1.2/1.3 200 mesh copper grids (Delta microscopies), which had been glow-discharged twice using a Pelco glow discharge system at 15 mA for 25 s. Samples were vitrified in 100% liquid ethane using a Mark IV Vitrobot (Thermo Fisher Scientific) by blotting for 3.5 s with Whatman No. 1 filter paper at 8°C and 100% relative humidity, after a 15 s waiting time.

### Cryo-EM data collection, processing, refinement and modeling

Data collection was performed on a Glacios transmission electron microscope (Thermo Fisher Scientific) operating at 200 kV, using the EPU automated image acquisition software (Thermo Fisher Scientific). Movies were collected on a Falcon 4i direct electron detector operating in counted mode at a nominal magnification of 240,000x (0.58 Å/pixel) using defocus range of −0.75 to −2.5 µm. Movies were collected over a 1.8-s exposure and a total dose of ∼40 e-/Å^2^.

The data processing workflow is summarized in **Figure S6**. Briefly, all movies were motion-corrected and dose-weighted with MotionCorr2^54^ and the aligned micrographs were used to estimate the defocus values with patchCTF within cryosparc^55^. CryoSPARC blob picker was used for automated particle picking. The resulting particles were extracted (binning 4, 2.32 Å/pixel) and used to obtain initial 2D classes. The classes without clear features or poorly aligned were discarded and the remaining particles were re-extracted (binning 2, 1.16 Å/pixel) and used for a new round of 2D classification. Then, an initial *ab initio* 3D model was generated in cryosparc and it was refined without imposing symmetry. After refinement, the particles were subjected to the cryosparc 3D classification into 10 classes without initial volumes or masks. The maps from the different classes were analyzed to assign them a spike conformation (closed, 1-RBD-up, 2-RBD-up, open, or unidentifiable). The map from each state that presented more density was selected for non-homogeneous refinement in cryoscparc (C3 symmetry applied only to the “closed” conformation map)^56^. The final map was sharpened with DeepEMhancer^57^ and the local resolution was estimated in cryosparc.

Model building started with an AlphaFold3 model of the trimeric full-length spike (ipTM: 0.82; pTM: 0.82), from which a single protomer was isolated and fitted into the sharpened EM map using UCSF Chimera. Then, regions connecting the individual domains were deleted, the separated domains were fitted individually, and the deleted regions were manually re-built in Coot. This initial model was refined with one round of real space refinement including morphing in Phenix^58^, followed by iterative rounds of manual building and real-space and B-factor refinement in Coot and Phenix, using secondary structure restraints. Then, symmetry operators were obtained from the EM map with the map-symmetry tool in Phenix, and they were used to place the three copies of the protomer within the trimeric map with the apply-NCS-operators Phenix tool. A final round of real-space refinement was performed in Phenix. Validation of model coordinates was performed using MolProbity.

### Biolayer interferometry (BLI)

The affinity of recombinant proteins was assessed in real-time using a bio-layer interferometry Octet-R8 device (Sartorius). Initial binding experiments were performed by loading nickel-nitriloacetic acid (Ni-NTA) capture sensors (Sartorius) for 10 min at 1,000 rpm shaking speed with the PHEV-RBD at 200 nM (or the SARS-CoV-2 RBD at 100 nM) in phosphate-buffered saline (PBS). The sensors were then blocked with PBS containing bovine serum albumin (BSA) at 1.0 mg/mL (assay buffer) and were incubated at 1,000 rpm with two-fold serially diluted concentrations (500 nM to 15.6 nM) of DPEP1 (or ACE2) in assay buffer. Association and dissociation were monitored for 240 s and 180 s, respectively. A sample reference measurement was recorded from a sensor loaded with each RBD and dipped in the assay buffer. Specific signals were calculated by subtracting nonspecific signals obtained for the sample reference from the signals recorded for the RBD-loaded sensors dipped in DPEP1 (or ACE2) solutions. Two independent experiments were performed but only the curves obtained at 500 nM in the first experiment were chosen for **Figure 3**.

Experiments to evaluate binding of the PHEV-spike were performed with DPEP1-Avi. This protein was biotinylated with a kit (Avidity) following the manufacturer’s instructions, and the reaction buffer was exchanged to PBS. Biotinylated DPEP1-Avi (50 nM dimer) was immobilized on streptavidin capture sensors (Sartorius) for 10 min at 1,000 rpm shaking speed. The sensors were then blocked with assay buffer and incubated with two-fold serially diluted concentrations (100 nM to 3.125 nM) of PHEV-spike (or Omicron BA.4/5 spike used as a control) in assay buffer. Association and dissociation were monitored for 240 s. A sample reference measurement was recorded from a sensor loaded with DPEP1-Avi and dipped in the assay buffer. After each dissociation step the sensors were regenerated by dipping them 30 s in acetate buffer (pH 4.0) and 30 s in PBS (three times).

Affinity of the recombinant PHEV-RBD towards DPEP1 was determined following a similar protocol. Streptavidin capture sensors (Sartorius) were loaded for 10 min at 1,000 rpm shaking speed with biotinylated DPEP1-Avi at 50 nM (dimer) in PBS. The sensors were then blocked with assay buffer, and they were incubated at 1,000 rpm with two-fold serially diluted concentrations (starting at 6,000 or 4,000 nM) of PHEV-RBD in assay buffer. Association and dissociation were monitored for 240 s. A sample reference measurement was recorded from a sensor loaded with DPEP1-Avi and dipped in the assay buffer. Regeneration steps were performed with acetate buffer (pH 4.0) as indicated before. The steady-state signal was plotted against the analyte concentration, and the curve was fitted assuming a 1:1 binding model. Six independent experiments were performed and the dissociation constant (*K_d_*) values from each of them were averaged and used to calculate the standard deviation.

### Statistical analysis

Statistics were performed in GraphPad Prism v10. All details about statistical tests can be found in the figure legends or in the main text.

## Acknowledgements

We thank the staff from the Crystallography platform at Institut Pasteur and the synchrotron source SOLEIL (Saint-Aubin, France) for granting access to the facility. We thank the staff of the beamline Proxima 2A for their advice and assistance during X-ray data collections. We also thank Pablo Guardado-Calvo for access to the Octet-R8. We acknowledge the NanoImaging Core at Institut Pasteur for support with sample preparation, image acquisition and analysis. The NanoImaging Core was created with the help of a grant from the French Government’s Investissements d’Avenir program (EQUIPEX CACSICE - Centre d’analyse de systèmes complexes dans les environnements complexes, ANR-11-EQPX-0008). This work was financially supported by an ERC Advanced Grant (101019724— EVADER) and a grant from the Spanish Ministerio de Ciencia e Innovación (PID2020-118602RB-I00— ZooVir) to R.S. J.D. is the recipient of an EMBO postdoctoral fellowship (ALTF-140-2021) and a Marie Skłodowska-Curie Actions Postdoctoral Fellowship (101104880).

## Authors’ contributions

J.D., I.F., F.A.R. and R.S. designed research; J.D., I.F., A.A. and A.H. performed research; J.D., I.F. and R.S. analyzed data; J.D., I.F., F.A.R. and R.S. wrote the manuscript; F.A.R. and R.S. provided funding. L.G.G.-L. provided essential material. All authors read and approved the final version of the manuscript.

## Data availability

All data supporting the findings of this study are available in this article. The structures of the PHEV RBD-DPEP1 complex and the close PHEV spike trimer have been deposited to the Protein Data Bank (PDB) under the accession codes 9H0B and 9H3J, respectively. The cryo-EM maps of the closed, 1-RBD-up, 2-RBD-up, and open PHEV spike trimers have been deposited to the Electron Microscopy Data Bank (EMDB) under accession codes EMD-51827, EMD-51844, EMD-51845 and EMD-51846, respectively.

