## Supplementary_Figs_Tables for "Dipeptidase 1 is a functional receptor for coronavirus PHEV"

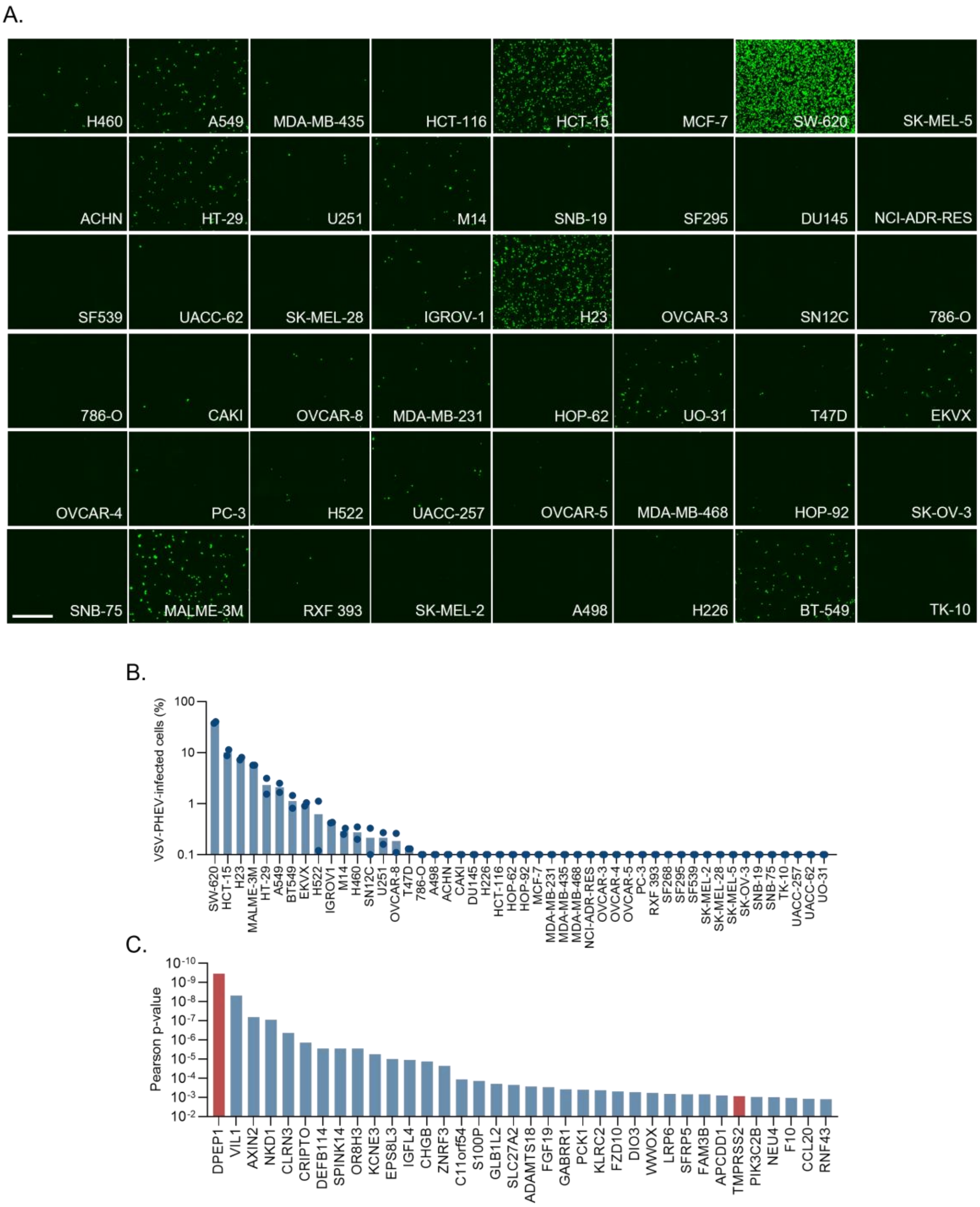


**Figure S1. Infection of the NCI-60 panel with PHEV pseudotypes.**

**A.** 48 cell lines from the NCI-60 panel were infected with PHEV pseudotypes and imaged 20 h later with the SX5 Live-Cell Analysis System at a 4X magnification. Images show the GFP signal. Cell line abbreviations are shown. Scale bar: 100 µm.

**B.** The percentage of infection of VSV-PHEV in the 48 cell lines of the NCI-60 panel is shown. The bar shows the mean and each dot represents an independent experiment (n = 2).

**C.** PHEV pseudotype infectivity was correlated with the expression of 7694 genes expressing membrane-associated proteins. The Pearson correlation p-values of the 40 genes best correlating with infectivity are shown.

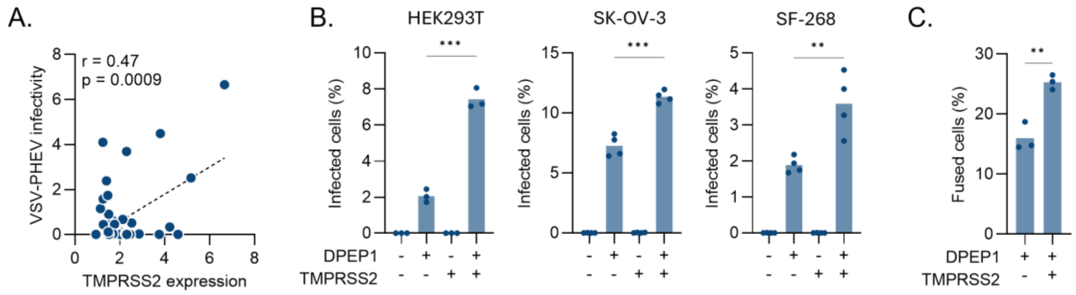


**Figure S2. TMPRSS2 enhances DPEP1-mediated PHEV entry**

**A.** Correlation between the expression of TMPRSS2 with the infectivity of PHEV pseudotypes in the NCI-60 cell line panel. Each point corresponds to a different cell line (n = 47). Pearsons r and p-value are indicated.

**B.** HEK293T, SK-OV-3 and SF-268 cells were transfected with DPEP1 and/or TMPRSS2 (or an empty vector) and infected with PHEV pseudotypes. The percentage of infected cells is shown.

**C.** HEK293T GFP-Split cells were transfected with DPEP1 and TMPRSS2 or an empty vector. The percentage of cell-cell fusion is shown.

In **B** and **C**, the bars show the mean, each dot represents a technical replicate (*n* = 3-4) from 1-2 independent experiments, and the significance of a two-sided unpaired t-test is indicated. ** P < 0.01; *** P < 0.005.

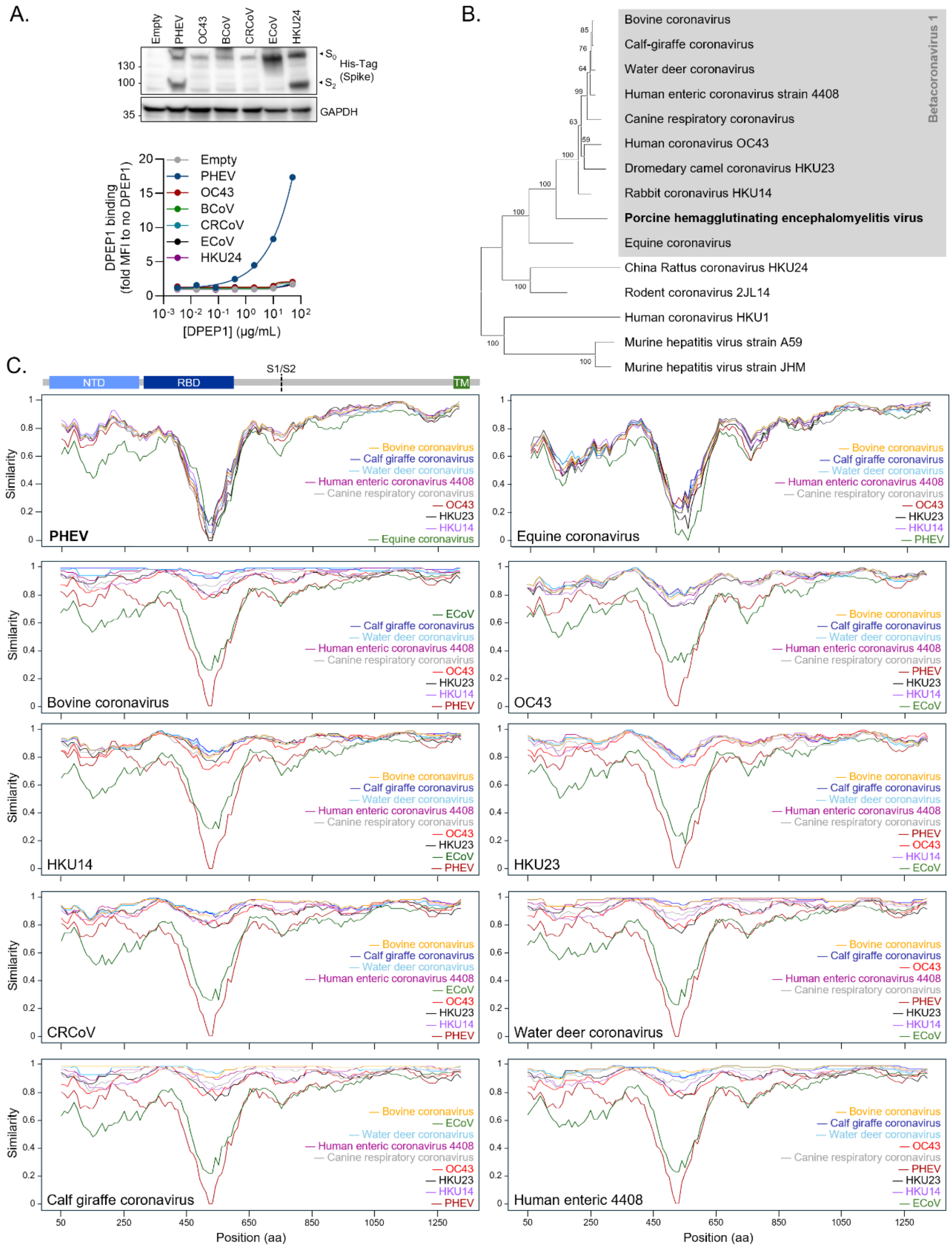


**Figure S3. DPEP1 is a PHEV-specific receptor.**

**A.** Flow cytometry measurement of soluble human DPEP1 binding to HEK293T cells transfected with betacoronavirus 1 spikes or an empty vector. Left: western blot confirmation of spike expression. A His-Tag added to the spike C-terminus was used for detection and GAPDH was used as a loading control. Right: the data shows the binding median fluorescence intensity (MFI) measured at each concentration of soluble DPEP1 normalized to the MFI in the absence of DPEP1, in cells transfected with an empty vector or the indicated betacoronavirus 1 spike. Results from a representative experiment are shown.

**B.** Maximum-likelihood phylogeny of embecovirus spike proteins. The grey shaded area shows the betacoronavirus 1 members. Bold font indicates the divergent Equine Coronavirus (ECoV) and Porcine Hemagglutinating Encephalomyelitis Virus (PHEV). Bootstrap values of each node are shown.

**C.** Similarity plots showing the percentage of similarity between the betacoronavirus 1 spikes. The spike used as a reference is indicated in the bottom-left corner. A schematic of the betacoronavirus 1 spike organization is shown at the top (NTD: N-terminal domain, RBD: receptor-binding domain, TM: transmembrane domain).

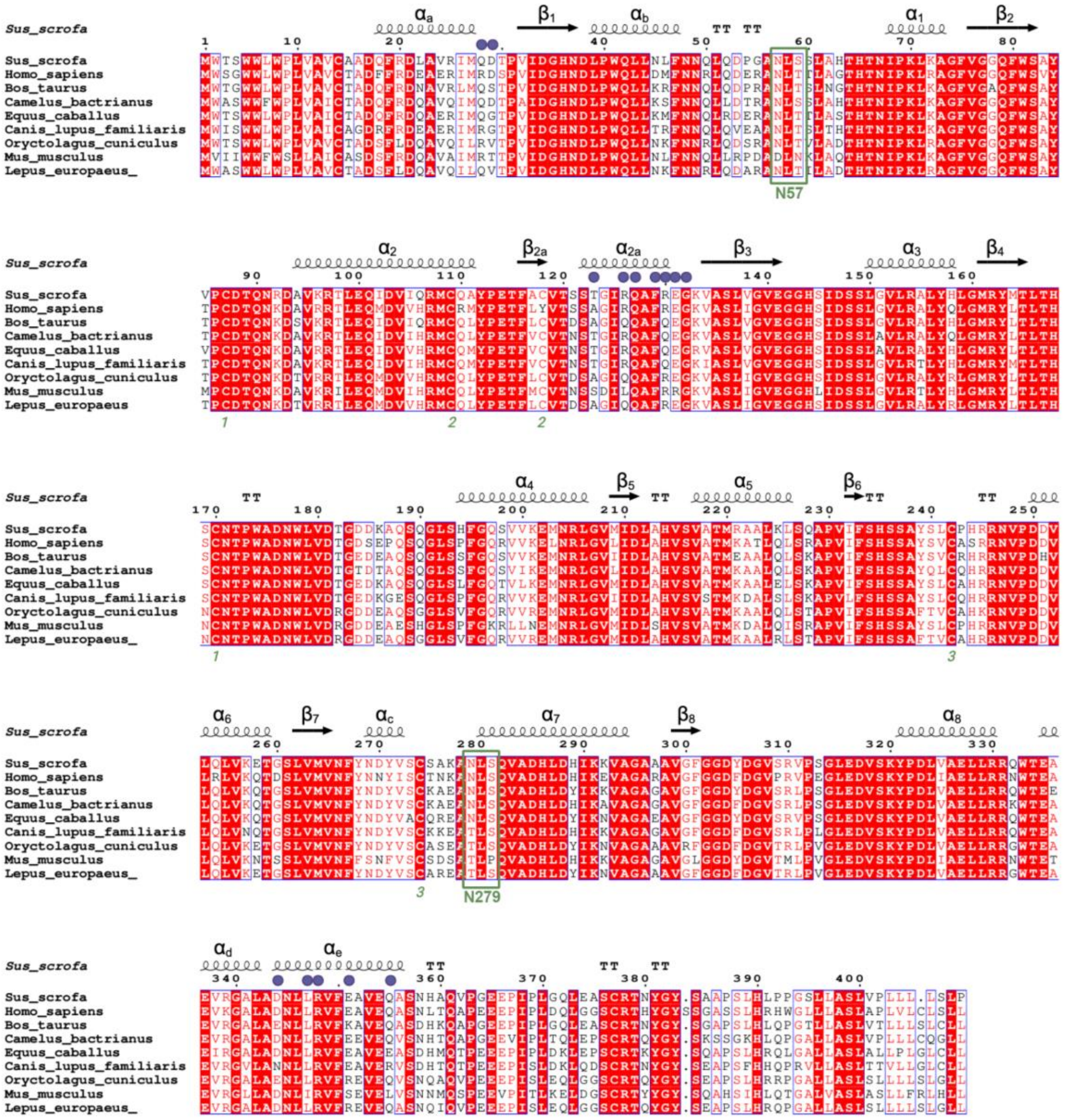

**Figure S4. The PHEV binding determinants on DPEP1 are partially conserved across mammals.** Amino acid sequence alignment of DPEP1 from selected mammals, calculated with MUSCLE[1] and rendered with ESPript 3.0[2]. Strictly conserved residues are in a red background and relatively conserved positions in red font. The secondary structure elements are labeled above the alignment. Residues contacted by the PHEV RBD are marked by violet full circles. Cysteines paired in disulfide bonds are indicated by the matching green numbers below the alignment. N-linked glycosylation sites are framed in green with the asparagine residue labeled. The sequences were obtained from the following accession numbers in Uniprot: *Sus scrofa*: P22412, *Homo sapiens*: P16444, *Bos taurus* Q3SZM7, *Camelus bactrianus*: A0A9W3FLA3, *Equus caballus*: F6TJM7, *Canis lupus familiaris*: A0A8P0THL7, *Oryctolagus cuniculus*: P31429, *Mus musculus*: P31428, *Lepus europaeus*: UPI002B47CA68.

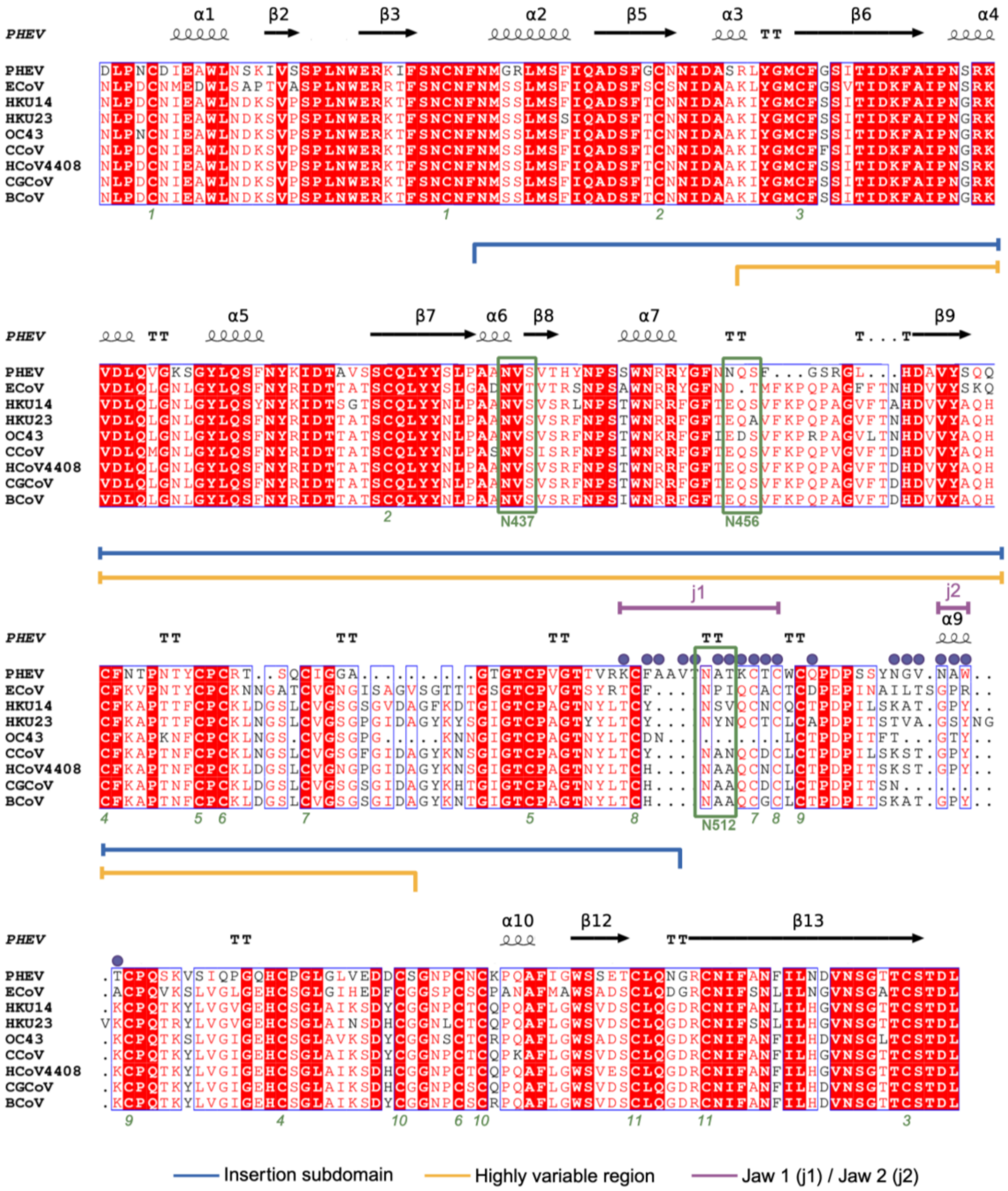


**Figure S5. Variable segments of the RBD recognize DPEP1.** Amino acid sequence alignment of the RBD region of the spike of members of the Betacoronavirus 1 species, calculated and rendered as in **Figure S4**, with small black dots indicating gaps. The secondary structure elements observed in the PHEV RBD are labeled above the alignment. The insertion subdomain is indicated with a blue line and the highly variable region (**Figure 1B**) by an orange line. Elements j1 and j2 introduced in **Figure 5** are in purple, and residues that contact DPEP1 in the structure are marked by violet full circles. Cysteines paired in disulfide bonds are indicated by the green number below the alignment. The three N-linked glycosylation sites in the PHEV RBD are framed in green (N437 conserved across the species, and N412 and N512 specific for PHEV). Note the important deletion in OC43 in the region of j1, which eliminates two cysteine residues involved in two different disulfide bonds. The crystal structure shows that the cysteines involved in disulfide bonds 7 and 8, left without their partner, form a disulfide bond with each other. The sequences used in the alignment were obtained from Uniprot with the following accession numbers: PHEV: A0A1V0IG63, ECoV: A8R4D0, HKU14: H9AA44, HKU23: X2JHN8, OC43: P36334, CCoV: Q7T5A1, HCoV4408: C6GHR0, CGCoV: A4ZU74, BCoV: Q9QAQ8.

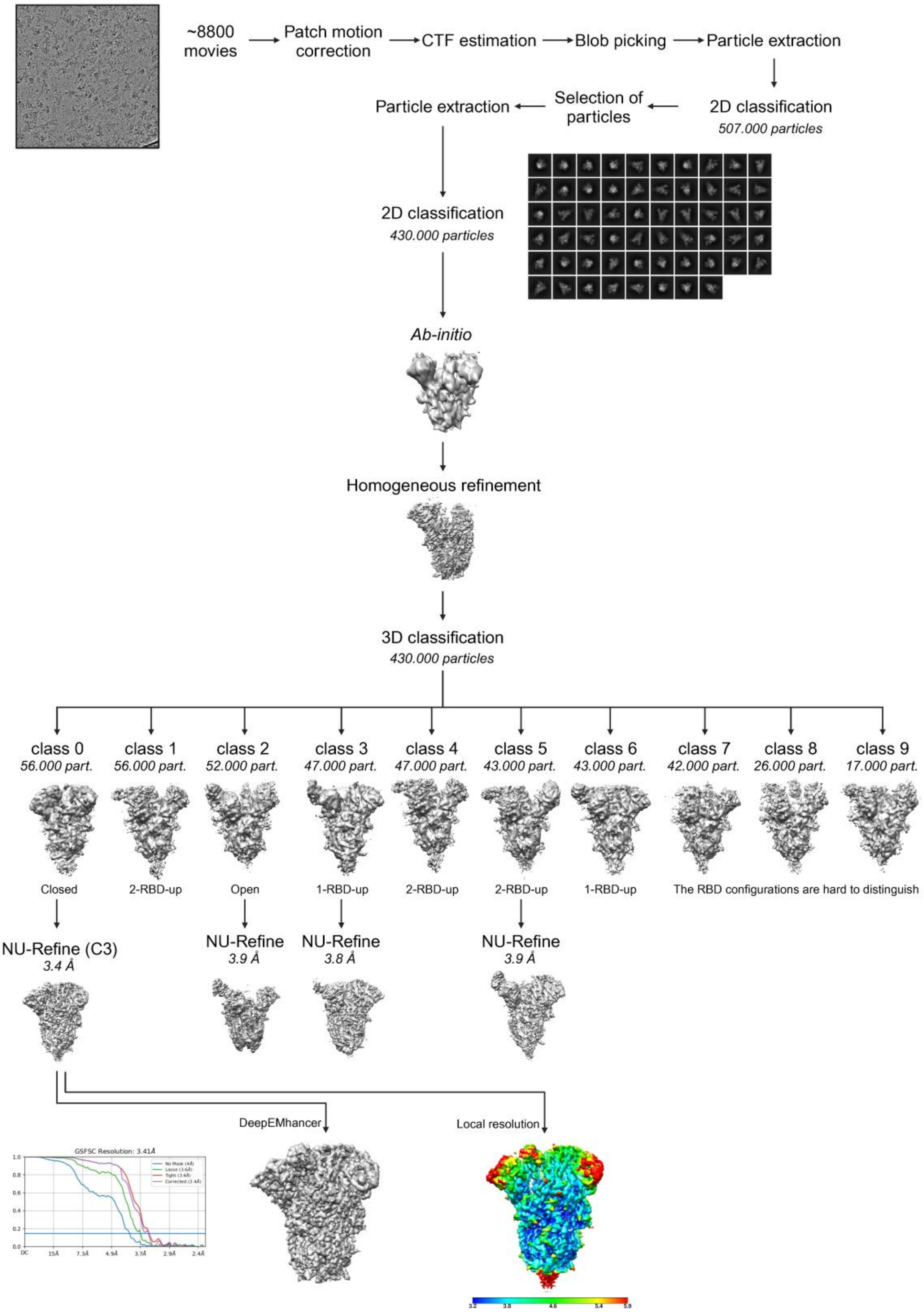


**Figure S6. Cryo-EM data processing workflow.**

Scheme detailing the steps followed to process the cryo-EM data collected for the PHEV spike ectodomain. A micrograph with particles, selected 2D class averages, the final sharpened map, a local resolution graphic, and the GSFSC (gold standard Fourier shell correlation) resolution plot are shown.

**Table S1. Data collection and refinement statistics**

| | PHEV-RBD + DPEP1 |
|---|---|
| PDB code | 9H0B |
| **Data Collection** | |
| Space Group | $P\ 1\ 2_1\ 1$ |
| a, b, c (Å) | 96.7, 73.0, 136.6 |
| a, b ,g (deg) | 90.0, 91.9, 90.0 |
| Resolution (Å) | 48.33-2.25 (2.31-2.25) |
| $R_{merge}$ | 0.238 (1.904) |
| Mean((I)/sd(I)) | 8.4 (1.0) |
| Number of unique reflections | 90,076 (6,350) |
| Completeness (%) | 99.6 (95.7) |
| Multiplicity | 6.9 (6.7) |
| $CC_{1/2}$ | 0.984 (0.454) |
| **Refinement** | |
| Number of reflections | 90,066 (2,715) |
| $R_{work}$ / $R_{free}$ | 19.87 / 23.55 |
| No. of atoms | |
| Macromolecules | 9,896 |
| Ligands | 224 |
| Solvent | 506 |
| Clashscore | 3.09 |
| R.m.s deviations | |
| Bond lenghts (Å) | 0.005 |
| Bond angles (°) | 0.760 |
| Ramachandran | |
| Favored (%) | 96.5 |
| Outliers (%) | 0.00 |
| Rotamer outliers | 0.73 |
| Average B-factor | 45.82 |
| Macromolecules | 45.25 |
| Ligands | 74.52 |
| Solvent | 44.20 |

Statistics for the highest-resolution shell are shown in parentheses.

**Table S2.** List of atoms involved in hydrogens bonds (cutoff 3.5 Å) and salt bridges (cutoff 4.0 Å) in the PHEV-RBD/DPEP1 complex (chains A and D/chains B and C, respectively).

| PHEV-RBD | DPEP1 | Distance in chains A-B (Å) | Distance in chains D-C (Å) |
|---|---|---|---|
| K505 (NZ) | D29 (OD1) | 3.75 | 2.80 |
| V530 (N) | R130 (O) | 3.04 | 3.06 |
| T514 (OG) | E351 (OE1) | 2.81 | 2.67 |
| T511 (OG1) | E351 (OE2) | 2.66 | 2.69 |
| C518 (O) | R130 (NH2) | 2.71 | 2.69 |
| Q521 (OE1) | R130 (NH2) | 3.06 | 3.04 |

**Table S3. Cryo-EM data collection, refinement, and validation statistics**

| | PHEV-spike ectodomain | | | |
|---|---|---|---|---|
| | **Closed conformation** | **1-RBD-up conformation** | **2-RBD-up conformation** | **Open conformation** |
| **PDB accession code** | 9H3J | | | |
| **EMDB accession code** | EMD-51827 | EMD-51844 | EMD-51845 | EMD-51846 |
| **Data collection and processing** | | | | |
| Magnification (x) | 240,000 | 240,000 | 240,000 | 240,000 |
| Voltage (kV) | 200 | 200 | 200 | 200 |
| Microscope | Glacios | Glacios | Glacios | Glacios |
| Camera | Falcon IV | Falcon IV | Falcon IV | Falcon IV |
| Electron exposure ($e^-$/Å$^2$) | 40 | 40 | 40 | 40 |
| Defocus range (μm) | -0.75 to -2.50 | -0.75 to -2.50 | -0.75 to -2.50 | -0.75 to -2.50 |
| Pixel size (Å) | 0.58 | 0.58 | 0.58 | 0.58 |
| Final particle images (number) | 55,920 | 47,399 | 43,088 | 52,409 |
| Symmetry imposed | C3 | C1 | C1 | C1 |
| Map resolution (Å) | 3.4 | 3.8 | 3.9 | 3.9 |
| FSC threshold | 0.143 | 0.143 | 0.143 | 0.143 |
| **Refinement and validation statistics** | | | | |
| Initial model | AlphaFold3 | | | |
| Model resolution (Å) | 3.5 | | | |
| Composition: | | | | |
| Chains | 6 | | | |
| Non-hydrogen atoms (number) | 23,358 | | | |
| Amino acids (number) | 2,940 | | | |
| Ligands (number) | NAG: 33 | | | |
| **Bonds (RMSD)** | | | | |
| Lengths (Å) [# > 4σ] | 0.003 | | | |
| Angles (°) [# > 4σ] | 0.624 | | | |
| **Ramachandran plot** | | | | |
| Favored (%) | 93.7 | | | |
| Allowed (%) | 6.3 | | | |
| Outliers (%) | 0.0 | | | |
| **Validation** | | | | |
| Rotamer outliers (%) | 3.2 | | | |
| C-β outliers (%) | 0.0 | | | |
| CABLAM outliers (%) | 3.9 | | | |
| CC (mask) | 0.76 | | | |
| Molprobity score | 2.4 | | | |
| Clash score | 11.4 | | | |

**Table S4. Primers used for PHEV spike and porcine DPEP1 mutagenesis**

| Gene | aa. change | Forward primer | Reverse primer |
|---|---|---|---|
| Pig DPEP1 | Q42A | gacctgccttggGCCctgctgaatctgttc | gaacagattcagcagGGCccaaggcaggtc |
| Pig DPEP1 | T123R | gcgtcaccagcagcaGaggcatccggcagg | cctgccggatgcctCtgctgctggtgacgc |
| Pig DPEP1 | R126A | gcagcacaggcatcGCCcaggccttccggg | cccggaaggcctgGGCgatgcctgtgctgc |
| Pig DPEP1 | E351A | ctgctgagggtcttcgCCgcagtggagcaggc | gcctgctccactgcGGcgaagaccctcagcag |
| Pig DPEP1 | E351R | ctgctgagggtcttcAGggcagtggagcaggc | gcctgctccactgccCTgaagaccctcagcag |
| Hare DPEP1 | R351E | gctgcgggtgttcGAggaggtagagcaagc | gcttgctctacctccTCgaacacccgcagc |
| Spike | Q487A | cttgccggacatccGCCtgcatcggcggagc | gctccgccgatgcaGGCggatgtccggcaag |
| Spike | K505A | caccaccgtgcgcGCCtgcttcgccgctgtg | cacagcggcgaagcaGGCgcgcacggtggtg |
| Spike | F507R | cgtgcgcaagtgcCGcgccgctgtgacaaac | gtttgtcacagcggcgCGgcacttgcgcacg |
| Spike | V510R | gtgcttcgccgctAGgacaaacgccacaaag | ctttgtggcgtttgtcCTagcggcgaagcac |
| Spike | T511D | gcttcgccgctgtgGACaacgccacaaagtg | cactttgtggcgttGTCcacagcggcgaagc |
| Spike | T514A | ctgtgacaaacgccGcaaagtgtacctgctg | cagcaggtacactttgCggcgtttgtcacag |
| Spike | T517R | cgccacaaagtgtaGAtgctggtgccagcc | ggctggcaccagcaTCtacactttgtggcg |
| Spike | Q521A | gtacctgctggtgcGCTcccgatcctagcac | gtgctaggatcgggAGCgcaccagcaggtac |
| Spike | K528E | gatcctagcacctacGagggcgtgaatgcc | ggcattcacgccctCgtaggtgctaggatc |
| Spike | N531A | cctacaagggcgtgGCtgcctggacctgcc | ggcaggtccaggcaGCcacgcccttgtagg |
| Spike | W533A | gggcgtgaatgccGCCacctgccctcagag | ctctgagggcaggtGGCggcattcacgccc |
